# Ultrafast molecular dynamics observed within a dense protein condensate

**DOI:** 10.1101/2022.12.12.520135

**Authors:** Nicola Galvanetto, Miloš T. Ivanović, Aritra Chowdhury, Andrea Sottini, Mark F. Nüesch, Daniel Nettels, Robert B. Best, Benjamin Schuler

## Abstract

Many biological macromolecules can phase-separate in the cell and form highly concentrated condensates. The mesoscopic dynamics of these assemblies have been widely characterized, but their behavior at the molecular scale has remained more elusive. Here we investigate condensates of two highly charged disordered human proteins as a characteristic example of liquid-liquid phase separation. The dense phase is 1000 times more concentrated and has 300 times higher bulk viscosity than the dilute phase. However, single-molecule spectroscopy in individual droplets reveals that the polypeptide chains are remarkably dynamic, with sub-microsecond reconfiguration times. We rationalize this behavior with large-scale all-atom molecular-dynamics simulations, which reveal an unexpectedly similar short-range molecular environment in the dense and dilute phases, suggesting that local biochemical processes and interactions can remain exceedingly rapid in phase-separated systems.

## Introduction

Biological macromolecules in the cell can form assemblies where high local concentrations of proteins and nucleic acids accumulate in phase-separated droplets or membraneless organelles (1-3). Phase separation plays a key role in cellular processes, such as ribosome assembly, RNA splicing, stress response, mitosis, and chromatin organization (4, 5), and is involved in a range of diseases (1, 3, 6). An essential driving force for such phase transitions is the multivalency of binding domains or motifs in the participating proteins. Such interactions are particularly prevalent for intrinsically disordered proteins (IDPs), which either lack a well-defined three-dimensional structure or contain large disordered regions that can mediate interactions with multiple binding partners (7-10). However, the dynamic structural disorder in these often liquid-like assemblies have rendered it challenging to obtain a molecular-scale investigation of their dynamical properties. NMR spectroscopy has provided evidence that IDPs in condensates can retain backbone dynamics on the pico-to nanosecond timescale and retain their disorder (3, 11), but most experimental information on condensate dynamics has been limited to translational diffusion and mesoscopic physical properties, such as viscosity and surface tension (12-14).

To extend our understanding beyond the mesoscopic level, we probe the dynamics at the molecular scale within a condensate using a combination of single-molecule spectroscopy and large-scale all-atom explicit-solvent molecular dynamics (MD) simulations. Single-molecule Förster resonance energy transfer (FRET) and nanosecond fluorescence correlation spectroscopy (nsFCS) allow us to obtain experimental information on intramolecular distance distributions on nanometer length scales and associated dynamics down to nanosecond timescales (15-18). MD simulations validated with such experimental data provide atomistic insight into the molecular conformations, dynamics, and interactions underlying the liquid-like properties of biomolecular condensates (9-11). An important step towards the feasibility of such simulations has been the development of more accurate biomolecular force fields that provide a realistic representation of IDPs (19-21).

Here we investigate condensates of two highly charged intrinsically disordered human proteins, histone H1 (net charge +53) and its nuclear chaperone, prothymosin α (ProTα, net charge -44). In dilute solution, these two IDPs form a dimer with picomolar affinity, although they fully retain their structural disorder, long-range flexibility, and highly dynamic character when bound to each other (22, 23) (Fig. 1A). Both proteins modulate chromatin condensation, they are involved in transcriptional regulation (4, 5, 24), and condensates of H1 are present in the nucleus (25). At high protein concentrations, solutions of ProTα and H1 exhibit phase separation into a protein-rich dense phase and a dilute phase. We find that despite the high viscosity of the dense phase, the individual IDPs retain rapid dynamics on the hundreds-of-nanoseconds timescale, surprisingly close to the behavior in the dilute phase. These rapid dynamics enable a direct comparison to large-scale MD simulations of ProTα-H1 condensates, which reveal the origin of the similarity: The electrostatic interactions between the IDPs are highly transient both in the dilute and the dense phase and involve a similar number of contacts on average. The resulting dynamic network reconciles slow translational diffusion with rapid chain dynamics, a behavior that may enable the occurrence of rapid local biochemical processes and interactions even in dense biomolecular condensates.

**Fig. 1.**
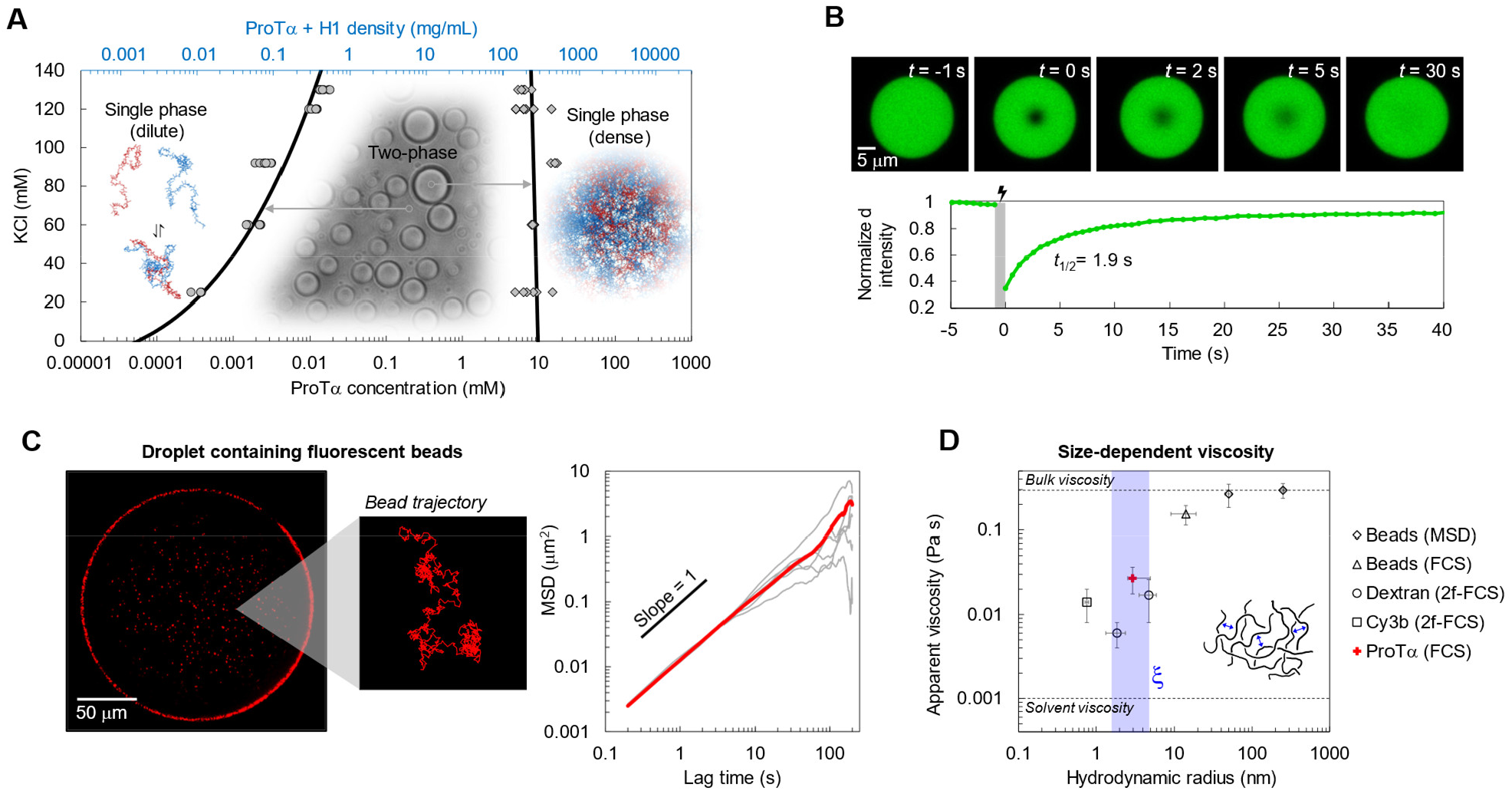
Mesoscopic properties of ProTα-H1 droplets. (**A**) Phase diagram as a function of salt concentration upon mixing ProTα and H1 at a 1.2:1 stoichiometric ratio. The total protein density (top axis) is based on the measured ProTα concentration (bottom axis) and charge balance in both phases (Fig. S1). Phenomenological fit (solid line) based on Voorn-Overbeek theory (31, 32). (**B**) Fluorescence recovery after photobleaching the center of a droplet doped with labeled ProTα at 120 mM KCl. (**C**) (left) Fluorescence image and representative trajectory of 500-nm beads diffusing in a droplet at 120 mM KCl. (right) Mean squared displacement (MSD) of five representative 500-nm beads (gray) and average (red). (**D**) Probe-size-dependent viscosity from measurements of diffusion coefficients for Cy3B, dextran, ProTα, and polystyrene beads in droplets at 120 mM KCl. Measurement of apparent viscosity by particle tracking (MSD, see **C**), single-focus fluorescence correlation spectroscopy (FCS), or two-focus FCS (2f-FCS). The shaded band indicates the correlation length, ξ, in the dense phase. See Methods for error bars and the estimate of ξ and the hydrodynamic radius of ProTα.

## Results

### ProTα and H1 phase-separate and form viscous droplets

The strong electrostatic interactions between ProTα and H1 (22, 26) can lead to co-phase separation, or complex coacervation, as observed for other highly charged biological and synthetic polyelectrolytes (8, 23, 27, 28). Especially at low salt concentrations, mixtures of the two proteins separate into two phases (Fig. 1A): a dilute phase with nanomolar to micromolar protein concentrations, where 1:1 complexes, i.e., heterodimers of ProTα and H1 dominate (22, 26) (Fig. S2), and droplets of a dense phase, with a total protein mass fraction of ∼20%, similar to other biomolecular condensates (29, 30). Since phase separation is strongest when ProTα and H1 are present at a ratio of 1.2:1, where their charges balance (Fig. S1), we investigated their phase behavior at this stoichiometry. The pronounced influence of the salt concentration that is evident from the phase diagram is reminiscent of the steep dependence of the binding affinity of the heterodimer on salt concentration (22, 26) and reflects the dominant role of Coulomb interactions between these highly charged IDPs. To probe the translational diffusion of protein molecules inside the droplets, we employed fluorescence recovery after photobleaching (FRAP) on a sample doped with nanomolar concentrations of fluorescently labeled ProTα. Bleaching with a small confocal laser spot in the dense phase results in rapid recovery within a few seconds (Fig. 1B), reflecting the liquid-like nature of the condensate.

To further characterize the solution properties of the dense phase, we used nanorheology based on the microscopic tracking of 500-nm fluorescent beads diffusing inside the droplets (Fig. 1C). From the mean-squared distance that the beads travel as a function of time, we obtained a viscosity of 0.30 ± 0.06 Pa s at 128 mM ionic strength according to the Stokes-Einstein relation (see Methods). The bulk viscosity of the ProTα-H1 coacervates is thus ∼300 times higher than that of water, and within the range of dense-phase viscosities previously observed for other biomolecular condensates (13, 14, 29, 33). Owing to the polymeric nature of the constituents, the dense-phase viscosity is expected to depend on the size of the probe used for monitoring translational diffusion (34). We thus employed probe particles with different hydrodynamic radii, from the fluorophore Cy3B and dextran of different molecular masses, whose diffusion we assessed with fluorescence correlation spectroscopy (FCS), to fluorescent beads of different radii (Fig. 1D). Across this size range, we indeed observed a pronounced increase in viscosity from ∼0.01 Pa s to ∼0.30 Pa s, with a transition near the correlation length, *ξ*, which is related to the effective mesh size of the underlying polymer network (13). The diffusion of molecules smaller than *ξ* is barely impeded by the mesh formed from the interacting IDP chains, whereas the motion of particles larger than *ξ* is strongly hindered and dominated by the bulk viscosity of the droplet. In summary, ProTα and H1 exhibit prototypical liquid-liquid phase separation with a dense-phase viscosity more than two orders of magnitude greater than that of the dilute phase. How is this large increase in viscosity linked to the structure and dynamics of the IDP chains making up the network?

### Observing rapid molecular dynamics in the dense phase

To investigate the behavior of individual protein molecules within the droplets, we doped the solution of unlabeled ProTα and H1 with picomolar concentrations of ProTα labeled with Cy3B as a FRET donor and CF660R as an acceptor at positions 56 and 110 (ProTαC). Confocal single-molecule FRET experiments allowed us to probe the conformations and dynamics of ProTα both in the dilute and in the dense phase (Fig. 2A–E). The mean FRET efficiency, ⟨*E*⟩, reports on intramolecular distances and distance distributions (18). Owing to efficient mutual screening of the two highly charged IDPs, ProTα is more compact when bound to H1 (⟨*E*⟩_*PH*_ = 0.55 ± 0.03) than in isolation (⟨*E*⟩_*P*_ = 0.35 ± 0.03) (22, 26) (Fig. 2F). As expected from the protein concentrations (26), the dimer is the dominant population in the dilute phase (Fig. S2). In the dense phase, we obtained values of ⟨*E*⟩ intermediate between these two values (Fig. 2F), indicating that ProTα is more expanded than in the dimer with H1, but more compact than in isolation.

**Fig. 2.**
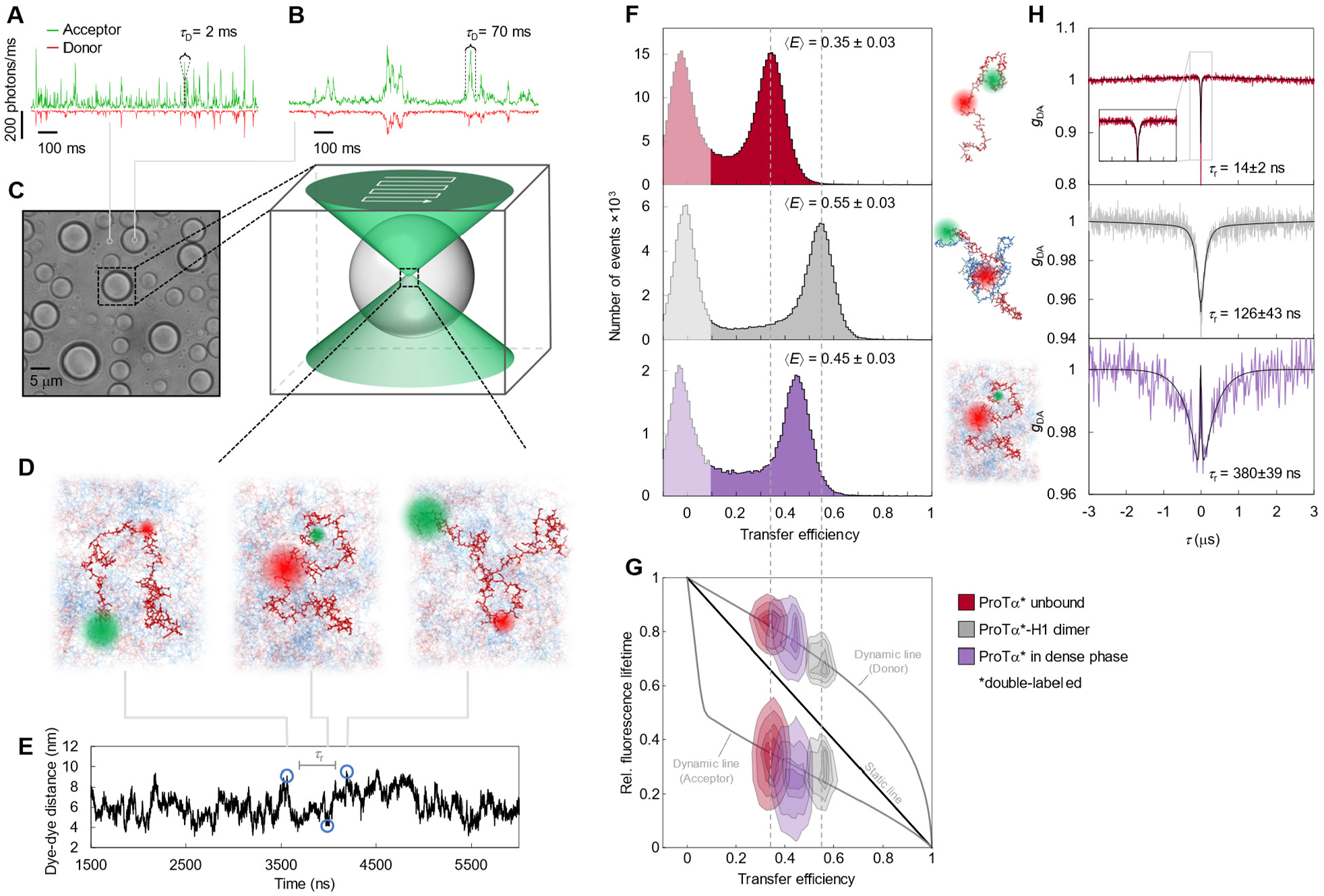
Single-molecule spectroscopy in the dilute and dense phases. (**A**) Photon time traces in the dilute phase (100 µW laser power) and (**B**) in the droplets (30 µW laser power in scanning mode, see **C**) with picomolar concentration of double-labeled ProTα. (**C**) Experimental scheme: single-molecule measurements were performed by positioning the confocal volume inside droplets that are stationary on the surface of the measurement chamber. (**D**) Configurations of double-labeled ProTα rapidly sampling different dye-dye distances and FRET-dependent fluorescence along a molecular trajectory from MD simulations (**E**). (**F**) Single-molecule transfer efficiency histograms of ProTαC (ProTα labeled at positions 56 and 110) in 128 mM ionic strength as a monomer in solution (top), in the heterodimer with H1 (middle), and within droplets (bottom) obtained with continuous-wave excitation while scanning at 3 μm/s in a serpentine pattern to improve statistics as shown in (**C**). (**G**) 2D histogram of relative donor and acceptor fluorescence lifetime versus FRET efficiency (18) for all detected bursts obtained with pulsed excitation. The straight line shows the dependence expected for fluorophores at a fixed distance; curved lines show the dependences for fluorophores exploring a distribution of distances (self-avoiding walk polymer, see Methods; upper line: donor lifetime; lower line: acceptor lifetime). (**H**) Nanosecond fluorescence correlation spectroscopy probing chain dynamics based on intramolecular FRET in double-labeled ProTα; data show donor–acceptor fluorescence cross-correlations with single-exponential fits (black lines) normalized to 1 at their respective values at 3 μs to facilitate direct comparison. Resulting reconfiguration times, τ_r_, are averages of three independent measurements (errors discussed in Methods).

The analysis of fluorescence lifetimes from time-correlated single-photon counting demonstrates the presence of broad distance distributions in all three cases (Fig. 2G), as expected for disordered proteins (18). Similar results were obtained for ProTα labeled at positions 2 and 56 (ProTαN, Fig. S3). Based on the single-molecule measurements, we infer average end-to-end distances (35) of 10.9 ± 0.5 nm, 9.2 ± 0.5 nm, and 9.4 ± 0.3 nm for ProTα alone, the dimer, and ProTα in the droplet, respectively (see Methods for details). The dimensions of ProTα in the droplet are in the same range as the correlation length in the dense phase (Fig. 1D), indicating that the proteins in the droplets are in the semidilute regime, where the chains can overlap but are not fully entangled (34). The expansion of ProTα relative to the dimer is thus suggestive of ProTα interacting with multiple H1 molecules simultaneously in the dense phase.

The broad intramolecular distance distributions entail the presence of highly heterogeneous conformational ensembles, but on which timescale do these conformations interconvert in the dense phase? We can probe these chain reconfiguration times, *τ*_r_, in single-molecule FRET experiments combined with nsFCS: Fluctuations in inter-dye distance cause fluctuations in the intensity of donor and acceptor emission, which can be quantified by correlating the fluorescence signal (18). Based on this approach, we measured *τ*_r_ = 14 ± 2 ns for unbound ProTα (36) and *τ*_r_ = 126 ± 43 ns in the ProTα-H1 dimer, as previously observed (22). To enable such measurements in the dense phase, we used longer-wavelength dyes compared to previous experiments (22) to reduce background from autofluorescence, and we combined nsFCS with sample scanning to compensate for the slow translational diffusion of the molecules in the droplets (Fig. 2H, Fig. S4). The resulting correlation functions yielded *τ*_r_ = 380 ± 39 ns, only a factor of ∼3 slower than the corresponding dynamics in the dimer, although the bulk viscosity of the droplets is ∼300 times greater than that of the dilute solution (Fig. 1D). Even if we consider the length scale dependence of viscosity (Fig. 1D), a large discrepancy remains between the relative slowdown of translation diffusion and chain dynamics. In summary, single-molecule FRET thus reveals a more expanded average conformation of disordered ProTα in the dense phase compared to the dimer and remarkably rapid intrachain dynamics. To elucidate the molecular origin of this behavior, we turned to MD simulations.

### Interaction dynamics from molecular simulations

Since we aim to compare absolute timescales with experiment, we require all-atom molecular dynamics simulations with explicit solvent. In view of the experimentally determined reconfiguration timescale of ∼380 ns for protein chains in the dense phase, a direct comparison is within reach. We thus performed large-scale simulations of a dense phase consisting of 96 ProTα and 80 H1 molecules (ensuring charge neutrality) in a slab configuration (37) in 128 mM KCl, corresponding to ∼4 million atoms in the simulation box (Fig. 3A). We employed the Amber ff99SBws force field (38) with the TIP4P/2005s water model (39), a combination that has previously performed well in IDP and condensate simulations (19, 37). Based on a total simulation time of ∼6 μs (Movie 1, Movie 2), and aided by the large number of protein copies in the system, we obtained enough sampling for a meaningful comparison with experimentally accessible quantities.

**Fig. 3.**
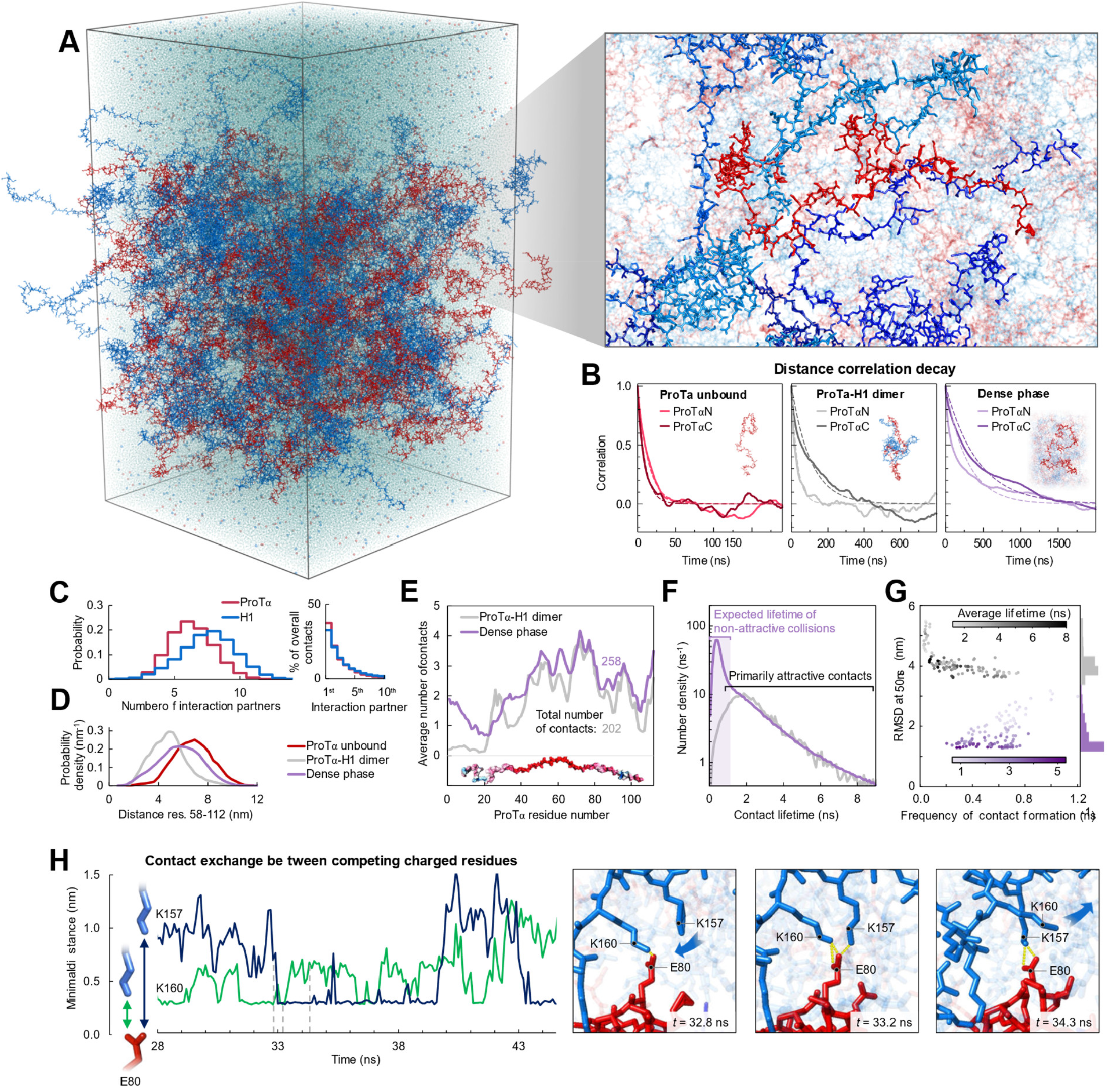
Large-scale molecular dynamics simulations of ProTα-H1 phase separation. (**A**) All-atom explicit solvent simulation system composed of 96 ProTα (red) and 80 H1 molecules (blue) in slab geometry (37), including water (light blue spheres), K^+^ ions (blue spheres), and Cl^-^ ions (red spheres). The zoom-in shows four H1 molecules (different shades of blue) interacting with one ProTα. See Movies 1–3 for an illustration of the dynamics. (**B**) Time correlation functions of the distance between residues 5 and 58 (ProTαN) and residues 58 and 112 (ProTαC) from simulations of ProTα unbound (left), in the heterodimer (middle), and in the dense phase (right), with single-exponential fits (dashed lines). (**C**) Histograms of the number of H1 chains simultaneously interacting with a single ProTα chain (red) and vice versa (blue). Contributions of each interaction partner to the total number of residue-residue contacts are shown on the right. (**D**) Distance distributions between ProTα residues 58 and 112 in the different conditions (see legend). (**E**) Average number of contacts that each residue of ProTα makes in the dimer (gray) and in the dense phase (purple), with average total number of contacts indicated. Only ∼11% of all contacts of ProTα in the dense phase are with other ProTα chains. (**F**) Distribution of the lifetimes of contacts formed by ProTα in the dimer (gray) and in the dense phase (purple). The areas under the curves correspond to the total number of new contacts formed per chain in one nanosecond. The shaded band indicates the contact lifetimes expected for non-attractive collisions (see Fig. S9 for details). (**G**) Root-mean-square displacement (RMSD) of the 112 individual ProTα residues within 50 ns vs their average frequency of contact formation. Average contact lifetimes are represented by the color scale; numbers of residues with similar RMSD are histogrammed on the right. (**H**) Example of rapid exchange between salt bridges in the dense phase, illustrated by two time trajectories of the minimum distance between the residue pairs involved (left) and corresponding snapshots from the simulation (right). See Movie 3 for an illustration of the rapid contact dynamics.

Both the total protein concentration and the translational diffusion coefficient of ProTα in the simulated slab are comparable to the experimental values (see Table 1) at the same salt concentration, suggesting that the viscosity and the overall balance of interactions in the simulations are realistic. Similarly, the average transfer efficiencies of ProTα from the simulations are close to the experimental values, both for ProTα in isolation, in the dimer, and in the dense phase (Fig. 2F, Table 1). Furthermore, the intramolecular distance distributions (Fig. 3D) are wide, as expected from the fluorescence lifetime analysis (Fig. 2G). Most remarkably, even the chain dynamics, based on intrachain distance correlation functions (Fig. 3B), are in the same range as the experimental result. Although the distribution of reconfiguration times, *τ*_r_, is broad owing to the remaining limitations of conformational sampling during the simulation time (40), the mean value of ∼400 ns compares well with experiment and is only a factor of ∼4 slower than in the dimer (Fig. 3B). Based on this validation by experiment, we scrutinize the simulations for the origin of such rapid chain dynamics despite the large viscosity in the dense phase.

**Table 1.** Comparison between observables from experiments (EXP) and simulations (MD) (⟨E⟩: average transfer efficiency; τ_r_: reconfiguration time). ProTαN and ProTαC refer to the measurements on the N-terminal and the C-terminal segment of full-length ProTα, respectively (see Table S1).

| Sample | Protein density<br>(mg/mL) | ProTα Diffusion coefficient<br>(m<sup>2</sup>/s) × 10<sup>-12</sup> | ProTαN |  | ProTαC |  |
| --- | --- | --- | --- | --- | --- | --- |
|  |  |  | ⟨E⟩ | τ<sub>r</sub> | ⟨E⟩ | τ<sub>r</sub> |
| ProTα [EXP] | — | 85 ± 9 | 0.41 ± 0.03 | 21 ± 2 ns | 0.35 ± 0.03 | 14 ± 2 ns |
| ProTα [MD] | — | 50 ± 7 | 0.49 ± 0.02 | 14 ± 4 ns | 0.30 ± 0.02 | 10 ± 3 ns |
| ProTα-1H dimer [EXP] | — | 74 ± 8 | 0.46 ± 0.03 | 64 ± 10 ns | 0.55 ± 0.03 | 0.13 ± 0.05 μs |
| ProTα-1H dimer [MD] | — | 32 ± 3 | 0.48 ± 0.08 | 32 ± 9 ns | 0.65 ± 0.07 | 0.11 ± 0.03 μs |
| Dense phase [EXP] | 220<sup>+210</sup><sub>-70</sub> | 2.7 ± 0.7 | 0.49 ± 0.03 | 0.30 ± 0.03 μs | 0.45 ± 0.03 | 0.38 ± 0.04 μs |
| Dense phase [MD] | 290 ± 10 | 1.5 ± 0.1 | 0.51 ± 0.11 | 0.29 ± 0.07 μs | 0.46 ± 0.18 | 0.4 ± 0.1 μs |

As expected from the optimal charge compensation between ProTα and H1 and the large protein concentration in the dense phase, with a mass fraction of ∼20% (Fig S6), ProTα and H1 engage in a network of interactions with oppositely charged chains. Each ProTα molecule, e.g., interacts on average with ∼6 H1 molecules simultaneously (Fig. 3C) and is slightly more expanded than in the dimer (Fig. 3D), in line with the measured FRET efficiencies (Fig. 2F). These intermolecular networking effects are expected to cause the high viscosity observed in the droplets (34) (Fig. 1D), but how can the *intra*molecular chain dynamics remain so rapid? An important clue comes from the inter-residue contact profiles, which reveal comparable interaction patterns in the heterodimer and in the dense phase (Fig. 3E), suggesting a remarkable similarity in the local environment experienced by the individual protein molecules. Indeed, the total number of contacts that a ProTα chain makes in the dense phase is only ∼28% greater than in the dimer, mainly due to contributions from the chain termini (Fig. 3E).

Another important insight comes from the lifetimes of these inter-chain contacts: In contrast to the persistent interactions expected for more specific binding sites (28), the duration of individual contacts between residues in ProTα and H1 is at most a few nanoseconds (Fig. 3F, Fig. S7, Fig. S8), orders of magnitude shorter than the chain reconfiguration time. Individual contacts thus never become rate-limiting for the motion of the polypeptide chain. The distributions of the longest contact lifetimes, above ∼2 ns, are very similar in the dimer and the dense phase, but a discrepancy is apparent for very short-lived contacts, which are much more prevalent in the dense phase (Fig. 3F). Many of these events can be attributed to the N-terminus of ProTα, whose fleeting encounters with other proteins in the crowded environment occur on a timescale expected for non-attractive random collisions (Fig. S9). Notably, this N-terminal region of ProTα makes hardly any contacts with H1 in the dimer because of its low net charge (22) (Fig. 3E). The lack of specific residue-residue interactions combined with the high concentrations of competing interaction partners in the dense phase can thus lead to rapid exchange between individual contacts (Fig. 3H, Fig. S12). It is worth emphasizing that the total concentration of charged side chains in the dense phase is in the range of 1 M.

Despite the similarity in the local environments and the kinetics of contact formation for the heterodimer and the dense phase, there are also notable differences. In contrast to the simple Brownian translational diffusion of the dimer in the dilute phase, individual proteins in the dense phase exhibit subdiffusion at timescales below the reconfiguration time (Fig. S10A), indicating locally correlated dynamics among polymers in the crowded semidilute regime (41). At the level of individual amino acid residues, we observe a broad distribution of mobilities (Fig. 3G), but on average, residues in the dimer are more mobile than those in the dense phase. Fig. 3G shows that among the residues in the dimer, those that make more contacts tend to be the less mobile, as expected. In the dense phase, however, we observe the opposite behavior, where higher mobility exhibits some correlation with a higher frequency of contact formation. These contacts are primarily due to the short-lived fleeting collisions of the N-terminal residues, suggesting that these contacts are a byproduct of the high protein density but hardly impede chain motion. In contrast, residues that experience more long-lived contacts exhibit lower mobility and pronounced subdiffusion (Fig. S11). Overall, subdiffusion is much more prominent in the dense phase than in the dimer (Fig. S10, Fig. S11), reflecting different dynamic regimes of contact formation and chain interactions in the two phases that lead to the remaining differences in reconfiguration times.

## Discussion

The combination of our experiments and simulations provides a comprehensive picture of the ProTα-H1 condensate across a wide range of length and timescales. At length scales much greater than the mesh size, the condensate appears as a continuous viscous medium, ∼300 times more viscous than water (Fig. 1D). At short length scales, the apparent viscosity within the polymer network is reduced (34), which facilitates rapid intramolecular dynamics. MD simulations validated by their agreement with the experimental data provide an unprecedented atomistic view of the condensate; they point to two main conclusions: As opposed to the dilute phase, which is dominated by one-to-one interactions between ProTα and H1, the dense phase is formed by an extended network of multivalent interactions between the oppositely charged proteins (Fig. 3C), which causes the large macroscopic viscosity (34). At the molecular scale, however, the picture is surprisingly dynamic: The dense phase is a semidilute unentangled solution in which the proteins remain highly solvated; they rearrange rapidly; and their contacts with other chains exchange quickly and are exceedingly short-lived compared to the global chain reconfiguration dynamics. The resulting average local environment that a protein experiences — within a Bjerrum length of about 1 nm — is strikingly similar in the dense and the dilute phases, and the average number of contacts that a residue makes is dominated by its charge (Fig. 3E).

The behavior we observe is a remarkable example of the subtle balance of intermolecular interactions in biomolecular phase separation: On the one hand, the interactions must be strong enough for the formation of stable condensates; on the other hand, they need to be sufficiently weak to allow translational diffusion and retain fluid-like dynamics within the dense phase and molecular exchange across the phase boundary — processes that are essential for function, such as biochemical reactions occurring in condensates (1-3, 42). Our results on the two nuclear IDPs ProTα and H1 indicate that charge-driven condensates — of which there are many in the nucleus — can enable surprisingly rapid dynamics on molecular length scales by facilitated breaking and forming of contacts. This highly dynamic regime of interactions may contribute to a faster local exploration of binding partners in condensates and efficient biochemical reactions. Similarly, the kinetics of molecular self-assembly processes that require large rearrangements of the chain, including the formation of amyloid-like structures within condensates (6, 43), may not be strongly hindered by the dense yet liquid-like environment.

The combination of single-molecule spectroscopy with all-atom molecular simulations is a promising strategy for probing the molecular dimensions and dynamics in condensates. The close agreement of our experimental results with the simulations indicates that current atomistic force fields are of suitable quality not only for describing isolated IDPs (19) but even for their complex multimolecular interactions in condensates (37). The chemical detail and timescales of dynamics available from such experimentally validated simulations are an ideal complement to computationally less demanding coarse-grained simulations (20, 22, 23), which have proven powerful for describing thermodynamic and structural aspects of biomolecular condensates (9, 19, 32). Single-molecule spectroscopy inside live cells (44) may enable intracellular measurements, e.g. in charge-driven biomolecular condensates in the nucleus (4, 5). We also note that in spite of a century of research on the complexation of synthetic polyelectrolytes (27, 28) and a more recent understanding of their parallels with disordered biomolecules (8, 26, 45), the dynamics of individual polymer chains in these systems have remained largely elusive. Our approach is likely to be transferrable to synthetic polymers, thus offering a general strategy for deciphering the molecular basis of such dense polymeric environments.

## Methods

### Protein preparation and labeling

Recombinant wild-type human histone H1.0 was used (H1; New England Biolabs M2501S). ProTαC and unlabeled ProTα were prepared as previously described (26); ProTαN cloned into a pBAD-Int-CBD-12His vector was prepared according to a previously described protocol (46). Cysteine residues introduced at positions 2 and 56, and 56 and 110, respectively, were used for labeling the protein with fluorescent dyes (see SI Table 1 for all protein sequences). Before labeling the double-Cys variants of ProTα, the proteins in phosphate-buffered saline (PBS), pH 7, 4 M guanidinium hydrochloride (GdmHCl), and 0.2 mM EDTA were reduced with 10 mM Tris(2-carboxyethyl)phosphine hydrochloride (TCEP) for one hour. Subsequently, the buffer was exchanged to PBS pH 7, 4 M GdmHCl, 0.2 TCEP, and 0.2 mM EDTA without TCEP via repeated (5x) buffer exchange using 3-kDa molecular weight cut off centrifugal concentrators. The protein variants were labeled with Cy3B maleimide (Invitrogen) and CF660R maleimide (Invitrogen) using a protein-to-dye ratio of 1:6:6, and incubated for one hour at room temperature and overnight at 277 K. The excess dye was quenched with 10 mM DTT for ten minutes and then removed using centrifugal concentrators. The labeled protein was purified by reversed-phase HPLC on a Reprosil Gold C18 column (Dr. Maisch, Germany) without separating labeling permutants. The correct masses of all labeled proteins were confirmed by electrospray ionization mass spectrometry.

### Turbidity measurements

Turbidity measurements were performed using a NanoDrop 2000 UV-Vis spectrophotometer (Thermo Scientific). ProTα was added to a fixed volume of an H1 solution to achieve a final concentration of 10 µM H1 and investigate a wide range of ProTα:H1 ratios. The samples were mixed by rapid pipetting for ∼10 s, and turbidity was measured at 350 nm. Four measurements were made for every sample in rapid succession, and the turbidity values averaged. Prior to mixing, the stocks of both proteins were diluted in identical buffers.

### Confocal single-molecule spectroscopy

For single-molecule measurements, concentration determination, and fluorescence correlation spectroscopy, we used a MicroTime 200 (PicoQuant) equipped with an objective (UPlanApo 60×/1.20-W; Olympus) mounted on a piezo stage (P-733.2 and PIFOC; Physik Instrumente GmbH), a 532-nm continuous-wave laser (LaserBoxx LBX-532-50-COL-PP; Oxxius), a 635-nm diode laser (LDH-D-C-635M; PicoQuant), and a supercontinuum fiber laser (EXW-12 SuperK Extreme, NKT Photonics). Florescence photons were separated from scattered laser light with a triple-band mirror (zt405/530/630rpc; Chroma), separated first into two channels with a polarizing or a 50/50 beam splitter and finally into four channels with a dichroic mirror to separate donor and acceptor emission (T635LPXR; Chroma). Donor emission was additionally filtered with an ET585/65m band-pass filter (Chroma) and acceptor emission with a LP647RU long-pass filter (Chroma), followed by detection with SPCM-AQRH-14-TR single-photon avalanche diodes (PerkinElmer).

### Protein concentration determination in the dilute and dense phases

For protein concentration measurements, a mixture of unlabeled proteins (12 μM ProTα and 10 μM H1, charge-balanced), doped with a small concentration (∼10 pM to 10 nM) (30) of labeled ProTα in TEK buffer (10 mM Tris-HCl, 0.1 mM EDTA, pH 7.4, and varying amounts of KCl) was allowed to phase-separate at 295 K. For measurements in the dilute phase, the phase-separated mixture was centrifuged at 295 K for 30 mins at 25,000 *g*, such that the dense phase coalesced into a one large droplet. The supernatant was carefully aspirated and transferred into plastic sample chambers (μ-Slide, ibidi) for microscopy. For measurements in the dense phase, the phase-separated mixture was directly transferred to the sample chambers, and droplets were allowed to settle on the bottom surface of the sample chamber by gravity; the boundaries of individual droplets were identified via 3D confocal imaging, and FCS and intensity measurements were performed inside the droplets. The fluorescent labels were excited with 635-nm continuous wave laser light at 5 μW (measured at the back aperture of the objective), and the fluorescence photons were separated with a polarizing beam splitter and recorded on two detectors. The total ProTα concentrations in the dense and the dilute phases were obtained by dividing the concentrations of labeled ProTα, measured using FCS or intensity detection (see below), with the known doping ratio. The doping ratio was chosen so that the fluorescence signal from labeled ProTα in the samples was within the linear detection range, which required higher doping ratios for dilute-phase compared to dense-phase measurements. For every condition measured, at least two estimates of concentrations were obtained, one from FCS and one from intensity measurements. In most cases, however, measurements were replicated several times, also with different doping ratios. As indicated by turbidity measurements, the maximum formation of dense phase occurs at a molar ProTα:H1 ratio of 1.2:1 (Fig. S1), corresponding to charge balance, so all experiments were performed by mixing the two proteins at this ratio, and H1 concentrations were inferred from the ProTα concentrations based on this ratio in both the dilute and the dense phases.

We employed both fluorescence correlation spectroscopy (FCS) and quantitative fluorescence intensity measurements on a MicroTime 200 (PicoQuant) to determine the concentrations of double-labeled ProTα (Cy3B and CF660R at residues 56 and 110) in the dense and dilute phases (30) by exciting CF660R with 635-nm laser light. Correlation functions were fitted with a model for translational diffusion through a 3D Gaussian-shaped confocal volume:

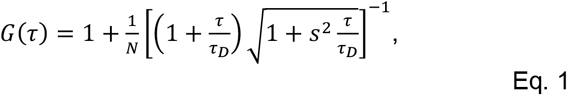

where *τ* is the lag time, *N* is the average number of fluorescent molecules in the confocal volume, *τ*_D_ is the translational diffusion time, and *s* is the ratio of the lateral and axial radii of the confocal volume. *N* is proportional to the concentration of labeled molecules, which can thus be estimated from FCS based on a calibration curve (30). The calibration curve was obtained by measuring samples of known concentrations of labeled ProTα (0.3-, 1-, 3-10-, 30-, and 100-nM) in buffer (10 mM Tris-HCl, 0.1 mM EDTA, pH 7.4, 120 mM KCl). The laser power used for the experiments and calibration was 5 μW (measured at the back aperture of the objective).

The average number of labeled proteins in the confocal volume *N* was also determined from the fluorescence intensity, and further corrected to yield *N*_*corr*_ = *N*(1 − *b*/*f*)^2^, as previously described (30), where *b* and *f* are the background and ProTα fluorescence intensities, respectively, and *N*_corr_ is used for concentration estimation. The background was estimated from samples without labeled protein. Similar to *N* obtained from FCS, the background-subtracted fluorescence intensity given by the mean photon count rates is proportional to protein concentration, and can thus also be used for concentration estimation based on the calibration curve.

### Fluorescence recovery after photobleaching (FRAP)

FRAP experiments were performed on a Leica SP8 confocal microscope with an HC PL APO CS2 63X/1.4 NA oil immersion objective. An area of ∼1.5 μm^2^ in droplets doped with ∼10 nM labeled ProTα (Cy3B and CF660R at residues 56 and 110) was bleached with a laser beam (530 nm wavelength) for 1 second, and fluorescence recovery was recorded by rapid confocal scanning. Images were processed with the Fiji open-source software (47), and recovery curves were analyzed in Mathematica (Wolfram Research) by fitting them with a single-exponential decay function. No aging or changes in the fluidity of the droplets were observed over the course of our observations (up to about four days).

### Nanorheology

We mixed 12 μM unlabeled ProTα and 10 μM unlabeled H1 with a small aliquot of fluorescent beads (100 nm and 500 nm diameter, Thermo Fisher Scientific) and centrifuged the sample to obtain a single droplet (diameter ≳100 μm). The motion of the beads was tracked at 295 K with an Olympus IXplore SpinSR10 microscope using a 100×/1.46 NA Plan-Apochromat oil immersion objective for 300 s with 50-mn exposure time and 200-ms time intervals. Trajectories were obtained with the ImageJ plugin TrackMate (48) and analyzed with custom MATLAB (MathWorks) code. Mean square displacements (*MSD*) were calculated in 2 dimensions and averaged (n = 22 for 100-nm beads, n = 20 for 500-nm beads). The diffusion coefficient, *D*, was calculated from

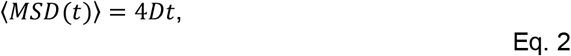

where *t* is the lag time. The viscosity was determined from the Stokes–Einstein equation assuming freely diffusing Brownian particles of radius *R* in a solution of viscosity *η*:

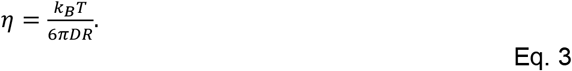

### Two-focus fluorescence correlation spectroscopy

Two-focus fluorescence correlation spectroscopy (49) measurements were performed at 295 K on a MicroTime 200 equipped with a differential interference contrast prism. Pulsed interleaved excitation with two orthogonally polarized supercontinuum fiber lasers (EXW-12 SuperK Extreme, NKT Photonics, equipped with a z520/5 band pass filter (Chroma), and Solea, PicoQuant, operating at 520 ± 3 nm) was used to form two laser foci. Both lasers were operated at a power of 5 μW (measured at the back aperture of the objective) and a repetition rate of 20-MHz, with the SuperK electronics triggering the Solea with a phase difference of half a period. The distance between the two foci was calibrated as previously described (50) with reference samples of Cy3b (51) and Dextran 10 kDa (52). The diffusion coefficient was determined by fitting the correlation functions as previously described (53) using Fretica (https://schuler.bioc.uzh.ch/programs).

### Hydrodynamic radii, apparent viscosity, and correlation length

Hydrodynamic radii (*R*_h_) of the beads were used as provided by the supplier. For dextran, we used the *R*_h_ values reported previously (52); we report the uncertainty based on the size-dependent polydispersity of our samples as estimated by the manufacturer. *R*_h_ of Cy3B was measured with two different techniques previously (51); we used the average value and provide the difference between the two as an uncertainty. *R*_h_ for a polymer diffusing in a semidilute solution is less well defined, so for ProTα we used a value for *R*_h_ extrapolated from experiments of ProTα in dilute solution. Based on the root-mean-square end-to-end distance (*r*_*rms*_) of ProTα measured in the dense phase (9.4 nm), we estimated the radius of gyration from *R*_g_ = *r*_*rms*_*/6*^1/2^. We observe the ratio *R*_g_/*R*_h_ for ProTα to be ∼1.3 in buffer, independent of salt concentration, so we used this ratio to obtain the corresponding value of *R*_h_ in the dense phase. As conservative estimates of uncertainty, we used as lower and upper bounds for this conversion the theoretical limits of *R*_g_/*R*_h_ for polymers (0.77 and 1.5) (50). Apparent viscosities were obtained from *D* and *R*_h_ using equation 3. Error bars of the apparent viscosity are the standard deviation of at least three measurements. The correlation length in the dense phase was estimated from *ξ* = *R*_g_ (*c*/*c*\*)^-3/4^, where *c* is the total protein concentration, and *c*\* is the overlap concentration (34). The range of *ξ* indicated as a shaded band in Fig. 1D was estimated by using either *R*_g_ or *R*_h_ for estimating *c*\* and by propagating the experimental error on *c* (Table 1).

### Single-molecule spectroscopy

Single-molecule measurements were performed at 295 K on a MicroTime 200 (PicoQuant). ProTα labeled with Cy3B and CF660R was excited with 532-nm continuous-wave laser light or in pulsed mode with alternating excitation of the dyes, achieved using pulsed interleaved excitation (54) at 20 MHz of the 635-nm diode laser and the SuperK supercontinuum fiber laser operated with a z532/3 band pass filter (Chroma). Measurements were performed in TEK buffer (10 mM Tris-HCl, 0.1 mM EDTA, pH 7.4, and 120 mM KCl). To avoid the pronounced adhesion of H1 to glass surfaces, plastic sample chambers (μ-Slide, ibidi) were used in all measurements. For single-molecule measurements in the dilute phase, the average power at the back aperture of the objective was 100 μW both for 532-nm continuous-wave excitation and for pulsed interleaved excitation (50 μW for donor and 50 μW for acceptor excitation), and the confocal volume was positioned 30 μm inside the sample chamber. FRET efficiency histograms in the dilute phase were acquired on samples with concentrations of labeled protein between 50 and 100 pM. For single-molecule measurements in the dense phase, the average power at the back aperture of the objective was between 10 and 30 μW both for 532-nm continuous-wave excitation and for pulsed interleaved excitation (5-15 μW for donor and 5-15 μW for acceptor excitation), depending on the background level, and the confocal volume was placed at the center of the spherical droplets, whose radius was between 4 and 15 μm. The samples were prepared by mixing unlabeled proteins (12 μM ProTα and 10 μM H1, charge balanced) with 5 to 10 pM of double-labeled ProTα.

Ratiometric transfer efficiencies were obtained from *E* = *n*_A_/(*n*_A_ + *n*_D_), where *n*_D_ and *n*_A_ are the numbers of donor and acceptor photons, respectively, in each burst, corrected for background, channel crosstalk, acceptor direct excitation, differences in quantum yields of the dyes, and detection efficiencies (55). From the transfer efficiency histograms, we obtained mean transfer efficiencies, ⟨*E*⟩, from fits with Gaussian peak functions. To infer end-to-end distance distributions, *P*(*r*), from ⟨*E*⟩, we use the relation (18)

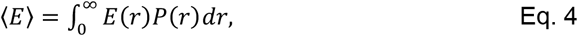

where

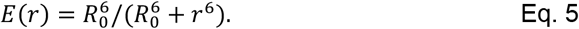

The Förster radius of 6.0 nm (56) for Cy3B/CF660R in water (*56*) was corrected for the refractive index, *n*, in the droplets. *n* increases linearly with the protein concentration (57) up to a mass fraction of at least 50 % (58). At 220 mg/mL, *n* is 3% greater than in water, resulting in *R*_0_ = 5.9 nm. For *P*(*r*), we applied an empirical modification of the self-avoiding-walk polymer model, the SAW-*v* model (35). We obtained the length scaling exponent, *v*, for the 2-56 and the 56-110 segments of ProTα, taking into account a total dye linker length for both fluorophores of 9 amino acids (59). To estimate the end-to-end distance of the complete ProTα chain, we used the total number of amino acids, *N*_*tot*_ = 110, and the average value of *v* obtained for the two segments. Note that fluorophore labeling has previously been shown to have only a small influence on the affinity between ProTα and H1 (22, 26). Since the fraction of labeled protein in the dense phase is only ∼10^−6^, a detectable effect of labeling on the dense-phase behavior is unlikely. Data analysis was performed using Fretica (https://schuler.bioc.uzh.ch/programs).

### Nanosecond fluorescence correlation spectroscopy (nsFCS)

Samples for nsFCS were prepared as described in the section *Single-molecule measurements*. To avoid signal loss from photobleaching in measurements inside droplets owing to the slow translational diffusion in the dense phase, the confocal volume (continuous-wave excitation at 532 nm) was continuously moved during data collection at a speed of 3 μm/s in a serpentine pattern (Fig. 2C) in the horizontal plane inside the droplet. Only photons from bursts of the FRET-active population (*E* > ⟨*E*⟩ − 0.15) were used for correlation analysis. Autocorrelation curves of acceptor and donor channels, and cross-correlation curves between acceptor and donor channels were computed from the measurements and analyzed as previously described (36, 40).

Full FCS curves with logarithmically spaced lag times ranging from nanoseconds to milliseconds are shown in Fig. S4. The equation used for fitting the correlations between detection channels *i, j* = *A, D* is

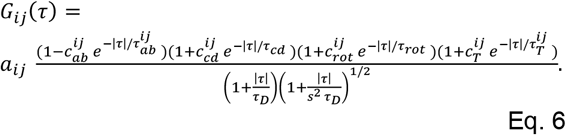

The four terms in the numerator with amplitudes *c*_ab_, *c*_cd,_ *c*_rot,_ *c*_T,_ and timescales *τ*_ab_, *τ*_cd,_ *τ*_rot,_ *τ*_T_ describe photon antibunching, conformational dynamics, dye rotation, and triplet blinking, respectively. *τ*_D_ and *s* are defined as in Eq. 1. Conformational dynamics result in a characteristic pattern with a positive amplitude in the autocorrelations (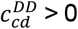 and 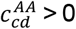) and a negative amplitude in the cross-correlation 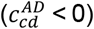, but with a common correlation time, *τ*_*cd*_. All three correlation curves (*G*_*DD*_(*τ*), *G*_*AA*_(*τ*), *G*_*AD*_(*τ*)) were fitted globally with *τ*_cd_ and *τ*_rot_ as shared fit parameters. *τ*_cd_ was converted to the reconfiguration time of the chain, *τ*_r_, as previously described (60), by assuming that chain dynamics can be modeled as a diffusive process in the potential of mean force derived from the sampled inter-dye distance distribution, *P*(*r*) (60, 61). The reported error of the reconfiguration time is either the standard deviation of three measurements or a systematic error of the fit, whichever was greater. The systematic error was estimated by fitting different intervals of the FCS data, especially by varying the lower bound of the fitted interval: We report as errors the range of reconfiguration times obtained by fitting from 0.8 ns and from 1.3 ns, a dominant source of variability in the results. We note that the conversion from *τ*_cd_ to *τ*_r_ does not entail a large change in timescale, and *τ*_cd_ and *τ*_r_ differ by less than 20% in all cases investigated here, depending on the average distance relative to the Förster radius (60).

There is evidence that rotation of the dyes is the origin of the correlated component at ∼30 ns given the asymmetry of the photon correlations when a polarizing beam splitter is used to separate the two major channels of detection (Fig. S5A-B). A similar timescale is present also in the relaxation of the intensity asymmetries of the parallel vs the perpendicular channel when the sample is illuminated with pulsed excitation (see Fig. S5C-D) (62).

### Fluorescence lifetime analysis

To get more information about the interdye distance distribution *P*(*r*), we determined in addition to *E* also the donor and acceptor fluorescence lifetimes, *τ*_*D*_ and *τ*_*A*_, for each burst. We first calculate the mean detection times, 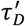 and 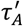, of all photons of a burst detected in the donor and acceptor channels, respectively. These times are measured relative to the preceding synchronization pulses of the laser pulsing electronics. Photons of orthogonal polarization with respect to the excitation polarization are weighted with 2*G* to correct for fluorescence anisotropy effects; corrects for the polarization-dependence of the detection efficiencies. For obtaining the mean fluorescence lifetimes, we further correct for the effect of background photons and for a time shift due to the instrument response function (IRF) with the formula: 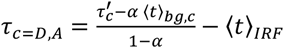, with α = *n*_*bg,c*_ Δ/*N*_*c*_. Here, ⟨*t*⟩_*bg,c*_ is the mean arrival time of background photons, ⟨*t*⟩_*IRF*_ is the mean time of the IRF, *n*_*bg,c*_ is the background rate, Δ the burst duration, and *N*_*c*_ is the total (uncorrected) number of photons in the donor (*c* = *D*) or acceptor (*c* = *A*) channels. The distribution of relative lifetimes, 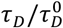 and 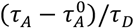, versus transfer efficiency are shown in Fig. 2G, where 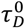 and 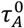 are the mean fluorescence lifetimes of donor and acceptor, respectively, in the absence of FRET. The theoretical dynamic FRET lines (63) in Fig. 2G were calculated assuming for *P*(*r*) the distance distribution expected from the SAW-*v* model (35). For the case that *P*(*r*) is sampled rapidly compared to the interphoton time (∼10 µs) but slowly compared to the lifetime of the excited state of the donor, it has been shown (64) that 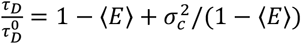 and 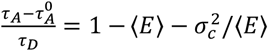, where the variance 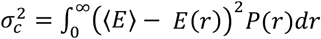. The dynamic FRET lines in Fig. 2G were obtained by varying the average end-to-end distance in the SAW-*v* model by changing *v*. The static FRET lines correspond to single fixed distances.

### Fluorescence anisotropy

We measured time-resolved fluorescence anisotropy decays (Fig. S5C,D) with pulsed interleaved excitation (54) of donor (Cy3B) and acceptor (CF660R) on double-labeled ProTα. We obtained time-correlated single-photon counting (TCSPC) histograms from photons polarized parallel and perpendicular with respect to the polarization of the excitation lasers. We corrected and combined them as previously described (65) to obtain the anisotropy decays for the acceptor (after direct acceptor excitation, Fig. S5D) and donor (after donor excitation, using donor-only bursts, Fig. S5C). Steady-state anisotropies of 0.18 for both Cy3B and CF660R indicate that rotational averaging of the fluorophores is sufficient for approximating the rotational factor *κ*^2^ by 2/3 (66).

### Molecular dynamics (MD) simulations

All-atom simulations of unbound ProTα, the ProTα-H1 dimer, and the phase-separated system were performed with the Amber99SBws force field (38, 67) with the TIP4P/2005s water model (38, 39). The temperature was kept constant at 295.15 K using stochastic velocity rescaling (68) (*τ* = 1 ps), and the pressure was kept at 1 bar with a Parrinello-Rahman barostat (69). Long-range electrostatic interactions were modeled using the particle-mesh Ewald method (70) with a grid spacing of 0.12 nm. Dispersion interactions and short-range repulsion were described by a Lennard-Jones potential with a cutoff at 0.9 nm. Bonds involving hydrogen atoms were constrained to their equilibrium lengths using the LINCS algorithm (71). Equations of motion were integrated with the leap-frog algorithm with a time step of 2 fs, with initial velocities taken from a Maxwell-Boltzmann distribution at 295.15 K. All simulations were performed using GROMACS (72), versions 2020.3 or 2021.5. We used the unlabeled variant of ProTα (Table S2) in all simulations, since the droplets under experimental conditions had 1000-fold higher concentration of unlabeled than labeled ProTα.

For the single ProTα chain, an initial extended structure was placed in a 20-nm truncated octahedral box. Subsequently, a short steepest descent minimization was performed, and the simulation box was filled with TIP4P/2005s water (38, 39) and again energy-minimized. In the next step, 518 potassium and 475 chloride ions (73) were added to the simulation box by replacing water molecules to match the ionic strength of the buffer used in the experiments (128 mM) and to ensure charge neutrality. Finally, a short energy minimization was performed for the whole system (809,843 atoms in total), before running molecular dynamics for a total simulation length of 3.19 µs. The first 100 ns were considered as system equilibration and were omitted from the analysis.

We performed 6 simulations of the ProTα-H1 dimer. The first four systems were constructed by placing extended ProTα and H1 chains close to each other (but not in contact, to minimize the initial structure bias) inside a 21-nm truncated octahedral box. Subsequently, the system was energy-minimized, and the simulation box was filled with TIP4P/2005s water (38, 39) and again energy-minimized. In the next step, 550 potassium and 560 chloride ions (73) were added to the simulation box by replacing water molecules to match the ionic strength of the buffer used in the experiment (128 mM) and to ensure charge neutrality. After the insertion of ions, the system (938,892 atoms in total) was again energy-minimized before initiating MD. The total simulation length of each of four runs was ∼3 µs. The first 300 ns of each run were considered as system equilibration and were omitted from the analysis. Runs 5 and 6 (∼2.2 µs each) were started from configurations at 1 µs of runs 1 and 2, respectively. The first 100 ns of runs 5 and 6 were omitted from the analysis to minimize the initial structure bias. In total, 15.15 µs of ProTα-H1 dimer simulations were used for the analysis.

The initial structure for all-atom simulations of the phase-separated system in slab configuration (37) was obtained with coarse-grained (CG) simulations. We utilized the one-bead-per-residue model that was previously developed to study the 1:1 ProTα-H1 dimer (22). Initially, 12 ProTα and 10 H1 molecules were randomly placed in a 25-nm cubic box, and the energy of the system was minimized with the steepest-descent algorithm. Although the CG model itself is capable of capturing the structure of the small globular domain (GD) of H1, we performed a 1-ns NVT run at 300 K with PLUMED (74) restraints, using the list of native contacts based on the experimental structure (75) (PDB 6HQ1), to ensure that the structure of the GDs was sufficiently close to the experimental structure (needed for all-atom reconstruction, below). In the next step, the box edge was decreased to 13.35 nm in a 30-ps NPT run to obtain an average protein density close to that of the dense phase in experiment. The system configuration was further randomized via a 280-ns NVT run at 500 K and an implicit ionic strength of 300 mM to ensure relatively uniform protein density in the box. Except for temperature and ionic strength, all other CG simulation parameters were identical to the parameters used in our previous study (22). Each chain from the final CG structure was independently reconstructed in all-atom form using a lookup table from fragments drawn from the PDB, as implemented in Pulchra (76). Side-chain clashes in the all-atom representation were eliminated via a short Monte Carlo simulation with CAMPARI (77) in which only the side chains were allowed to move. The relaxed configuration obtained with CAMPARI was multiplied 8 times, which, by tiling the box in X, Y, and Z directions, resulted in a 26.7-nm cubic box that contained 96 ProTα and 80 H1 molecules. Subsequently, the box edge was extended to 44 nm in the Z direction, and the resulting system was energy-minimized with the steepest-descent algorithm. To eliminate any non-proline cis-bonds that might have emerged during all-atom reconstruction, we ran a short simulation in vacuum with periodic boundaries, using a version of the force field that strongly favors trans peptide bonds (37) and applying weak position restraints to the protein backbone atoms and dihedral angles (5 kJ/mol/rad).

Subsequently, the simulation box was filled with TIP4P/2005s water (38, 39) and energy-minimized. In the next step, 2418 potassium and 2530 chloride ions (73) were added to the simulation box (4’000’932 atoms in total) to match the ionic strength of the buffer used in the experiments (128 mM) and to ensure charge neutrality. In the next step, the system was again energy-minimized, and a 20-ns MD run was performed with strong position restraints on protein backbone atoms (10^5^ kJ mol^-1^ nm^-2^) to stabilize any newly-formed trans bonds. Subsequently, a 1.7-ns simulation with PLUMED restraints on the native contacts of the GDs was performed to ensure that the structure of the reconstructed GDs was not perturbed during the equilibration procedure. The final structure of the run with native-contact restraints was used for the production run (with no restraints used), using GROMACS (72), versions 2020.3 and 2021.5. The free production run was 6.02 µs long, with a timestep of 2 fs. The first 1.5 µs were considered as system equilibration (Fig. S14) and were not used for the analysis.

### Analysis of MD simulations

Mean FRET efficiencies, ⟨*E*⟩, were obtained for each ProTα chain by calculating the instantaneous transfer efficiencies with the Förster equation (eq. 5) every 10 ps for both the ProTα-H1 dimer and the single ProTα chain simulations, and every 50 ps for every ProTα molecule in the dense-phase simulation. Subsequently, the instantaneous transfer efficiencies were averaged for each ProTα chain over the simulation length. Finally, ⟨*E*⟩ for the dimer was determined by averaging the transfer efficiencies calculated from six simulation runs, and ⟨*E*⟩ for the dense phase was determined by averaging over the 96 transfer efficiencies calculated for the individual chains. *R*_0_ = 6.0 nm was used for simulations of unbound ProTα and the ProTα-H1 dimer, *R*_0_ = 5.9 nm for the dense-phase simulations. Since we simulated ProTα without explicit representation of the fluorophores, the interdye distance, *r*, was estimated from the simulations via *r* = *d* ((*N* + 9)⁄*N*)^*v*^, where *d* denotes the distance between the Cα atoms of the labeled residues (residues 5-58 in ProTαN and residues 58-112 in ProTαC); *N* denotes the sequence separation of the labeling sites; and the scaling exponent *v* was set to 0.6 – we thus approximate the length of dyes and linkers by adding nine additional effective residues (78). The uncertainty in the transfer efficiency of the unbound ProTα chain was estimated from block analysis: the trajectory was divided into 3 intervals of equal length for which transfer efficiencies were calculated separately; the error was then taken as *σ*/√3, where *σ* is the standard deviation of these efficiencies. For the ProTα-H1 dimer, the transfer efficiency of ProTα was calculated as the average of the transfer efficiencies from six independent runs, and the error was estimated as the standard deviation. The transfer efficiency of ProTα in the dense-phase simulation was calculated by averaging the transfer efficiencies of 96 chains, and the error was estimated as the standard deviation of the average transfer efficiencies for the individual chains.

Chain reconfiguration times were estimated by integrating the residue-residue distance autocorrelations, *C*(*t*) (normalized to *C*(0) = 1), up to the time where *C*(*t*) = 0.03 and assuming the remaining decay to be single-exponential (79). For the simulation of unbound ProTα, the errors of the reconfiguration times were estimated by block analysis. For the ProTα-H1 dimer, autocorrelation functions from six independent simulations were determined, the reconfiguration times of ProTα chain were determined by analyzing the corresponding correlation functions as described above, and errors were estimated by bootstrapping: the data were randomly resampled 100 times with replacement, and the error was taken as the standard deviation of the correlation times obtained. In the dense-phase simulation, some chains sampled a relatively narrow range of distance values. To address this simulation imperfection, we omitted from the analysis those chains whose variance of transfer efficiency was below 0.05 (for ProTαN, 3 out of 96 chains were omitted; for ProTαC, 23 chains were omitted). The global mean and variance of the remaining chains were used to compute the correlation function, rather than the mean and variance for each run separately. Errors were estimated by bootstrapping from the set of reconfiguration times of the individual chains, using 200 samples with replacements per observable, similar to the procedure for the dimer.

The average number of H1 chains that simultaneously interact with a single ProTα chain, as well as the average number of ProTα chains that simultaneously interact with a single H1 chain (Fig. 3C) in the dense-phase simulation were determined by calculating the minimum distance between each ProTα and each H1 for each simulation snapshot. The two molecules were considered to be in contact if the minimum distance between any two of their Cα atoms was within 1 nm. The same contact definition was employed when calculating residue-residue contacts (Fig. 3E): Two residues were considered to be in contact if the distance between any of their Cα atoms was within 1 nm.

Lifetimes of residue-residue contacts were calculated by a transition-based or core-state approach (80). In short, rather than using a single distance cut-off to separate bound versus unbound states – which tends to underestimate contact lifetimes – separate cut-offs were used to determine the formation and breaking of contacts. For each pair of residues, a contact was based on the shortest distance between any pair of heavy atoms, one from each residue. Starting from an unformed contact, contact formation was defined to occur when this distance next dropped below 0.38 nm. An existing contact was considered to remain formed until the distance increased to more than 0.8 nm. Given the large number of possible contacts in the dense-phase simulation (342,997,336), the simulation was broken down into nine 500-ns blocks and each analyzed separately with parallelized code. Average lifetimes of each residue-residue contact were calculated by dividing the total bound time by the total number of contact breaking events for that contact. Intra-chain contacts were omitted from the analysis. Average lifetimes of each pair of ProTα-H1 residues (averaged over the different combinations of ProTα, H1 chains the two residues could come from) were calculated by dividing the total contact time (summed over all combinations of ProTα and H1 chains) of a specific residue pair by the total number of the contact breaking events for the same residues (summed over the same combinations of chains). Similarly, to calculate average lifetimes of residue-residue contacts according to the residue type (Fig. S8), we first identified all contacts involving a particular pair of residue types, in which one residue was from the ProTα chain, while the second one was from either H1 or ProTα. Subsequently, the average lifetime of that residue-residue combination was calculated by dividing the total bound time by the total number of contact breaking events for the contacts involving those residue types. Excess populations of specific residue-residue type pairs (Fig. S8 C,D) were determined by dividing the observed average number of contacts for a pair of residue types by the value that would be expected if residues paired randomly in a mean field approximation. The average number of contacts for a pair of residue types was calculated as a sum of all times that residues of those types were in contact, divided by the simulation length. The expected average number of contacts between two residue types (type 1 and 2) were calculated as *N f(1) f(2)*, where *N* is the average total number of contacts, and *f(1)* and *f(2)* are the fraction of residues of type 1 and 2, respectively.

The mean square displacement (MSD) of individual residues and of the center of mass (COM) of ProTα molecules were calculated using the Gromacs function *gmx msd*. For the ProTα-H1 dimer simulations, MSD curves of specific ProTα residues (residues 1 to 112) were averaged over six simulation runs. MSD curves of each ProTα residue for each of the 96 chains were calculated in four 1-µs blocks. Subsequently, MSD curves of each specific residue were averaged over all chains and blocks. The translational diffusion coefficient, *D*, of the COM of unbound ProTα was calculated by fitting the MSD with MSD(*t*) = 6*Dt* up to 700 ns, and the error was estimated from block analysis: MSD was calculated from each third of the trajectory (each part being ∼1 µs long); diffusion coefficients of each segment were determined by fitting them up to 250 ns, and the uncertainty given is the standard error of the mean. Diffusion coefficients of the COM of ProTα in the ProTα-H1 dimer were calculated by fitting the averaged MSD curves up to 1 µs, and the uncertainty was estimated as the standard error of the mean of the fits of six individual chains up to 500 ns. The diffusion coefficient of the COM of ProTα in the dense-phase simulation was calculated by fitting the MSD curve averaged over all 96 molecules up to 1 µs, and the uncertainty was estimated as the standard error of the mean of the fits of 96 individual chains. Diffusion exponents, *α*, for the diffusion of individual residues (Fig S11) were estimated by fitting their MSD with MSD(*t*) = 6*Dt*^*α*^ up to 2 ns, a range where the MSD curves are linear in double-logarithmic plots (Fig. S10).

Densities of protein, water, and ions from dense-phase simulations were calculated perpendicular to the longest slab axis (Z axis in Fig. S6), using the calculated density profiles between 15 nm and 30 nm (Fig. S6).

## Acknowledgements

We thank Alessandro Borgia, Madeleine Borgia, Hagen Hofmann, Rohit Pappu, and Andrea Soranno for helpful discussions, Ruijing Zhu and Pawe*ł Ł*ukija*ń*czuk for technical assistance in protein preparation, and Edward Lemke for providing the pBAD-Int-CBD-12His plasmid. This work was supported by the Swiss National Science Foundation (B.S.), the Novo Nordisk Foundation Challenge program REPIN (#NNF18OC0033926, B.S.), the Intramural Research Program of the National Institute of Diabetes and Digestive and Kidney Diseases at the National Institutes of Health (R.B.B.), the Forschungskredit of the University of Zurich (N.G.), and the European Union’s Horizon 2020 research and innovation programme under the Marie Sk*ł*odowska-Curie grant agreement ID 898228 (A.C.). We utilized the computational resources of Piz Daint and Eiger at the CSCS Swiss National Supercomputing Centre and of the National Institutes of Health HPC Biowulf cluster (http://hpc.nih.gov). Mass spectrometry was performed at the Functional Genomics Center Zurich. FRAP and bead tracking were performed with support of the Center for Microscopy and Image Analysis, University of Zurich.

