## Supporting Material for "Ultrafast molecular dynamics observed within a dense protein condensate"

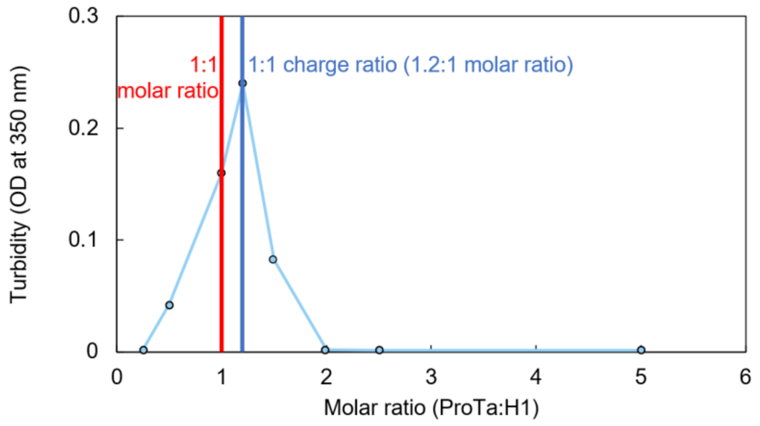


**Figure S1. Phase separation is most pronounced in a charged-balanced mixture of H1 and ProTα.** The extent of droplet formation was assessed using turbidity at a constant concentration of 10 μM H1 and varying amounts of ProTα at 50 mM KCl. Maximal phase separation was observed at a stoichiometric ratio of 1.2:1 for ProTα:H1, where the charges of the two proteins balance.

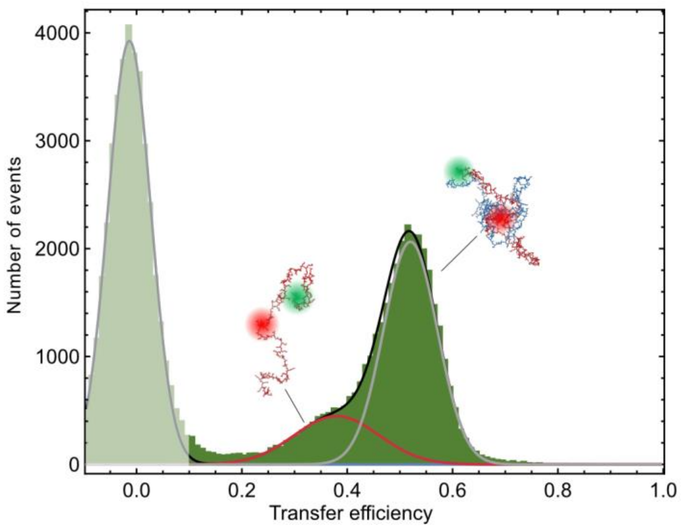


**Figure S2. The ProTα-H1 dimer is the dominant population in the dilute phase.** Single-molecule transfer efficiency histogram of ProTαC (labeled at position 56 and 110) in the dilute phase at 128 mM ionic strength. The phase-separated mixture was centrifuged, so that the dense phase coalesced into a single large droplet and no small droplets were diffusing in the dilute phase. The dilute phase was aspirated and transferred into a sample chamber for single-molecule measurements. In the fit (lines), the centers of the Gaussian peak functions were constrained to the transfer efficiencies measured with unbound ProTα and the ProTα-H1 dimer (Fig. 2F) to within experimental uncertainty. The shaded peak originates from molecules lacking an active acceptor dye.

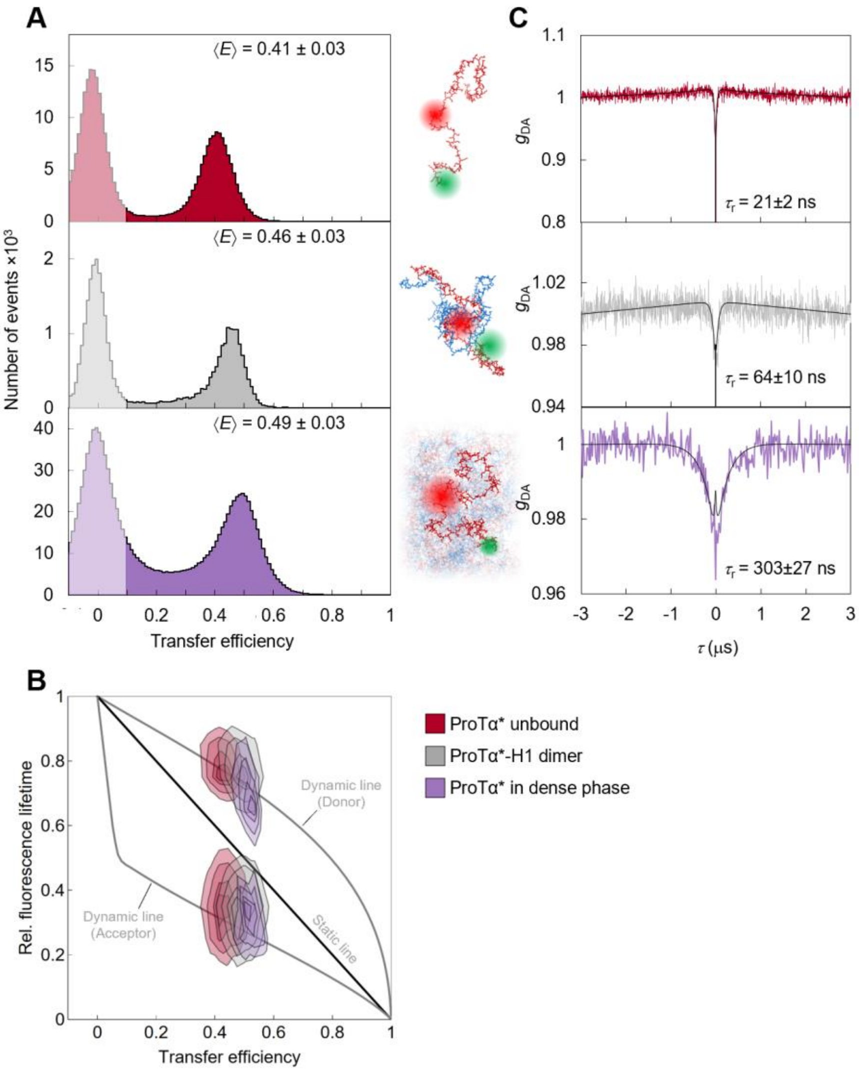


**Figure S3. ProTα labeled at positions 2 and 56 (ProTαN) shows behavior similar to ProTα labeled at positions 56 and 110 (ProTαC, Fig. 2).** (A) Single-molecule transfer efficiency histograms of ProTα (labeled with donor and acceptor at positions 2 and 56) at 128 mM ionic strength as a monomer in solution (top), in the 1:1 complex with H1 (middle), and within droplets (bottom) measured with continuous-wave excitation. Note the greater compaction in the dense phase compared to the ProTα-H1 dimer than for ProTαC. (B) 2D histogram of relative donor and acceptor fluorescence lifetimes versus FRET efficiency for all detected bursts measured with pulsed excitation. The straight line shows the dependence expected for fluorophores separated by a static distance; curved lines show the dependences for fluorophores that rapidly sample a distribution of distances (self-avoiding walk (SAWv (1)), see Methods; upper line: donor lifetime; lower line: acceptor lifetime). (C) nsFCS probing chain dynamics based on intramolecular FRET in double-labeled ProTαN; data show donor–acceptor fluorescence cross-correlations with fits (black lines). Reconfiguration times, $\tau_r$, are averages of $n = 3$ independent measurements (errors discussed in Methods).

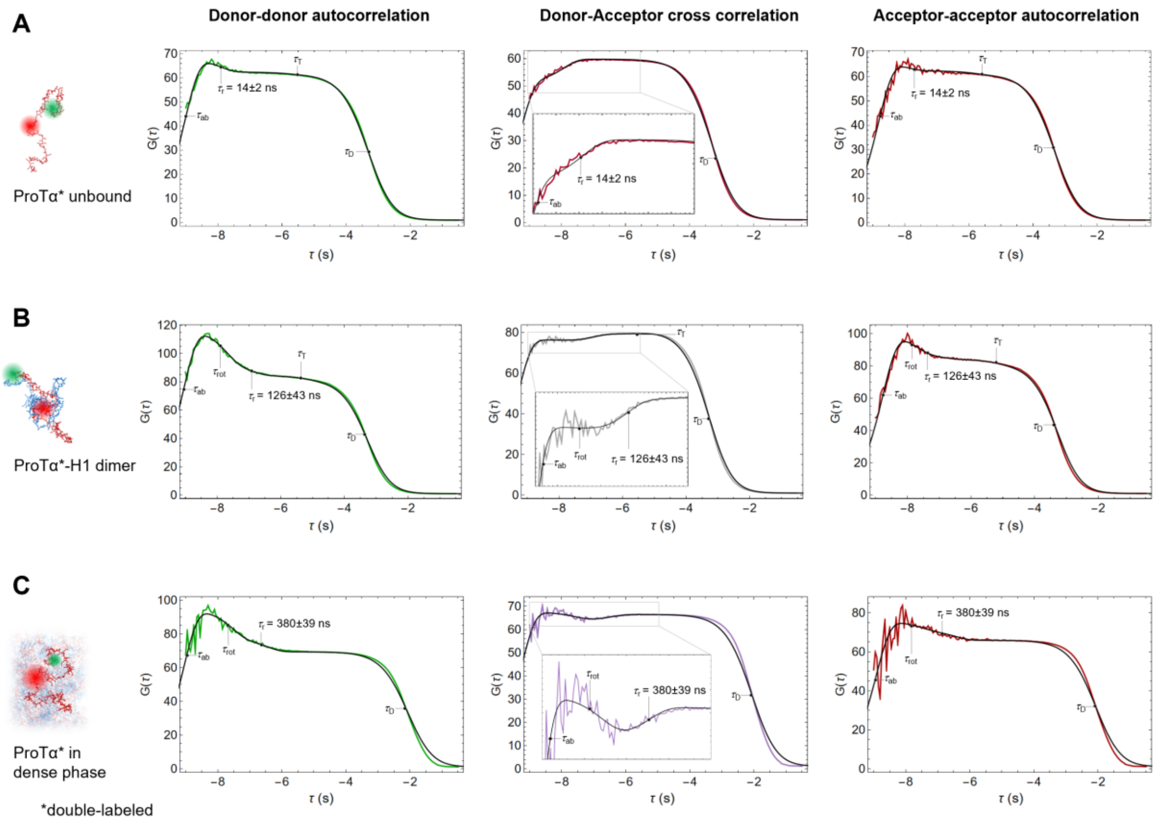


**Figure S4. FCS curves with logarithmic binning.** Donor and acceptor autocorrelations (green, red) and donor-acceptor crosscorrelations (purple; same color scheme as in Fig. 2H, which shows the same data with the same fits but in linear scale and normalized to an amplitude of 1 at ±3 μs) of ProTαC (labeled at position 56 and 110) in 128 mM ionic strength as an unbound monomer in solution (A), in the 1:1 complex with H1 (B), and within ProTα-H1 droplets (C). For each sample, the three correlations are fitted globally (see Methods; black solid lines) with shared correlation times for translational diffusion ($\tau_D$), triplet blinking ($\tau_T$), dye rotation ($\tau_{rot}$), and conformational dynamics ($\tau_{cd}$); photon antibunching ($\tau_{ab}$) is fitted individually. $\tau_{cd}$ was then converted to the reconfiguration time of the chain, $\tau_r$, as previously described (2) (we note that the conversion from $\tau_{cd}$ to $\tau_r$ does not entail a large change in timescale, and $\tau_{cd}$ and $\tau_r$ differ by less than 20% in all cases investigated here). $\tau_D$, $\tau_T$, $\tau_{rot}$, $\tau_r$, and $\tau_{ab}$ are shown in the panels when used in the fit, and they point to their corresponding timescales. The value of $\tau_r$ reported here is the mean of three measurements, as in Fig. 2H, and corresponds to the distance correlation time between the dyes at position 56 and 110 (2). $\tau_T$ in the donor-acceptor cross correlation in (B) shows a small negative amplitude, possibly indicating a slight contribution of slower distance dynamics on the microsecond timescale. Note the deviation between fit and measurement in (C) for the translational diffusion component is caused by focus scanning, which was required to improve statistics.

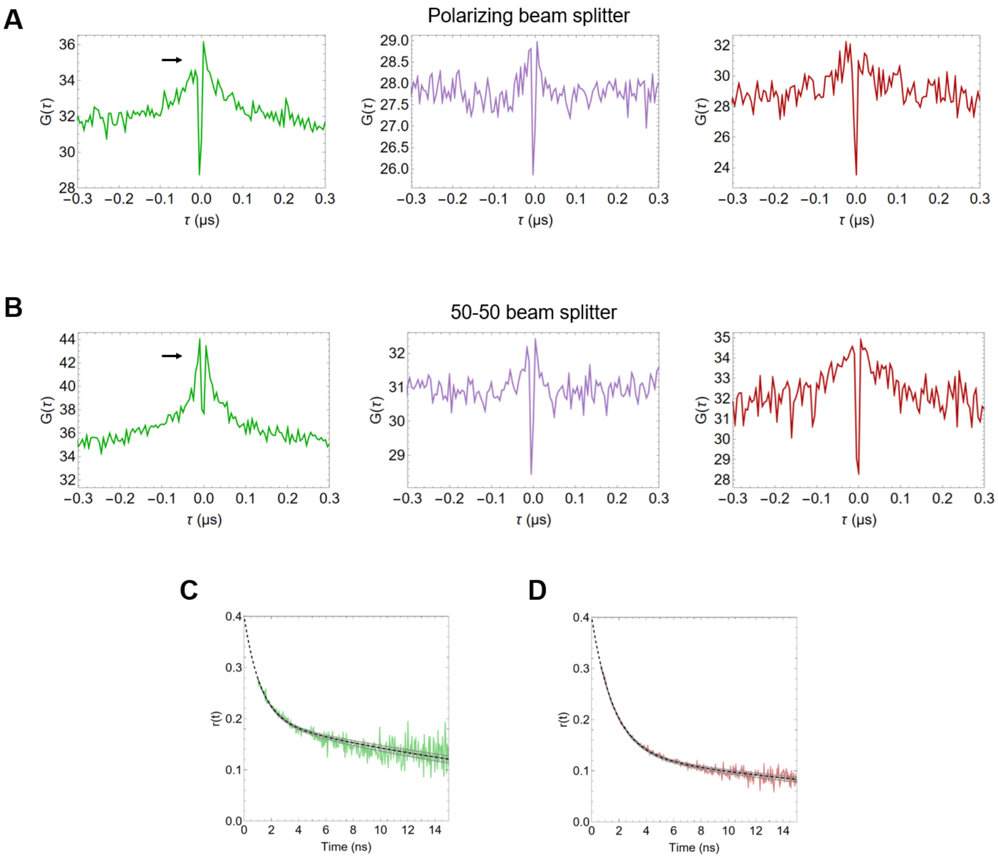


**Figure S5. Polarization-resolved fluorescence suggests that residual rotational dynamics in the dense phase causes correlated component in nsFCS.** (A) Donor and acceptor emission autocorrelations (green and red, respectively; parallel vs perpendicular channels) and donor-acceptor crosscorrelation (purple; sum of parallel and perpendicular channels) of the FRET-active subpopulation of labeled ProTαC in the dense phase when a polarizing beam splitter is used show asymmetry between the positive and negative lag-time branches in the positively correlated component (correlation time of 30 ns). This component is symmetric when a 50-50 beam splitter is used (B), indicating that the component is caused by polarization anisotropy. (C, D) Time-resolved anisotropy decays, *r*(*t*), from double-labeled ProTαC measured in the dense phase with pulsed interleaved excitation using (C) photons from donor-only bursts (transfer efficiency < 0.1, excitation at 532 nm) or (D) acceptor photons from bursts with transfer efficiency > 0.2 (excitation at at 635 nm). Data were fit with the function $r(t) = r_0((1 - A_{rot})e^{-\frac{t}{\tau_{fast}}} + A_{rot})e^{-\frac{t}{\tau_{rot}}}$ (black lines), with $r_0$ = 0.4 and the decay time $\tau_{\mathrm{rot}}$ = 30 ns obtained from the correlated component of the nsFCS (A, B) as a fixed fit parameter for the slow component of the decay. The agreement of the decay time of the slow component in *r*(*t*) with the decay time of the correlated component in the nsFCS further supports the role of residual rotation as the source of the latter.

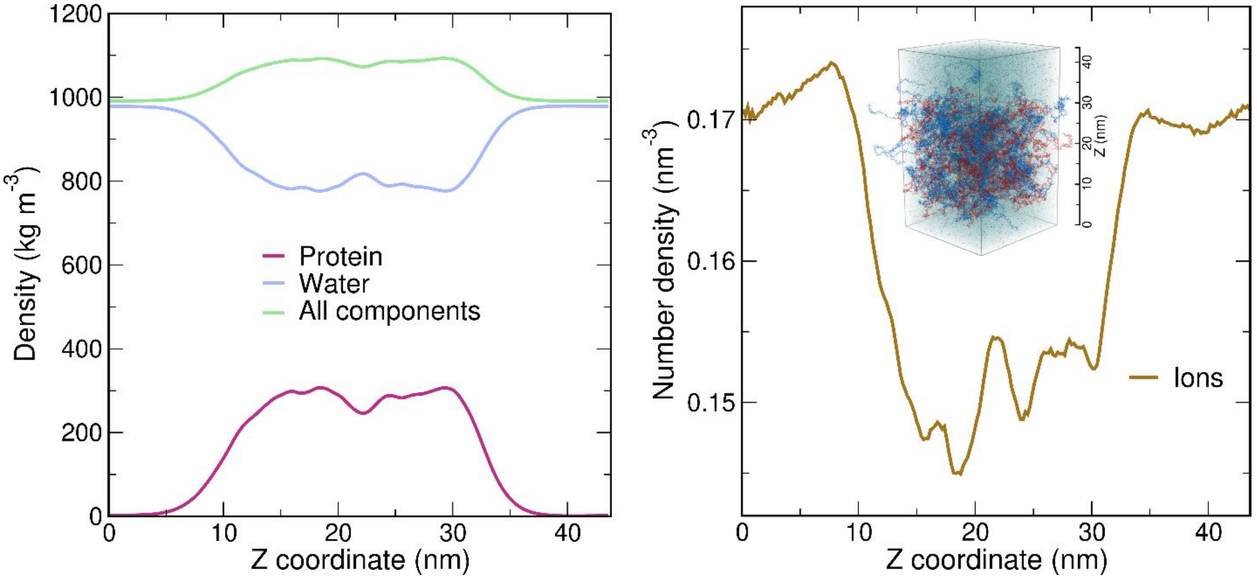


**Figure S6. Density profiles** of protein, water, all components (protein, water, and ions; left), and ions (right) along the Z axis (see inset on the right). The water density in the dense phase (central part of the slab, 15 nm < Z < 30 nm) is ~80.7% of the water density in the bulk regions (Z < 2.5 nm and Z > 40.0 nm). The number density of ions in the dense phase (15 nm < Z < 30 nm) is ~88.4% of the value close to the box edges (Z < 1.5 nm and Z > 41.0 nm). With respect to only the water density in the respective phases, the ion concentration is ~10% higher in the dense phase than in the dilute phase.

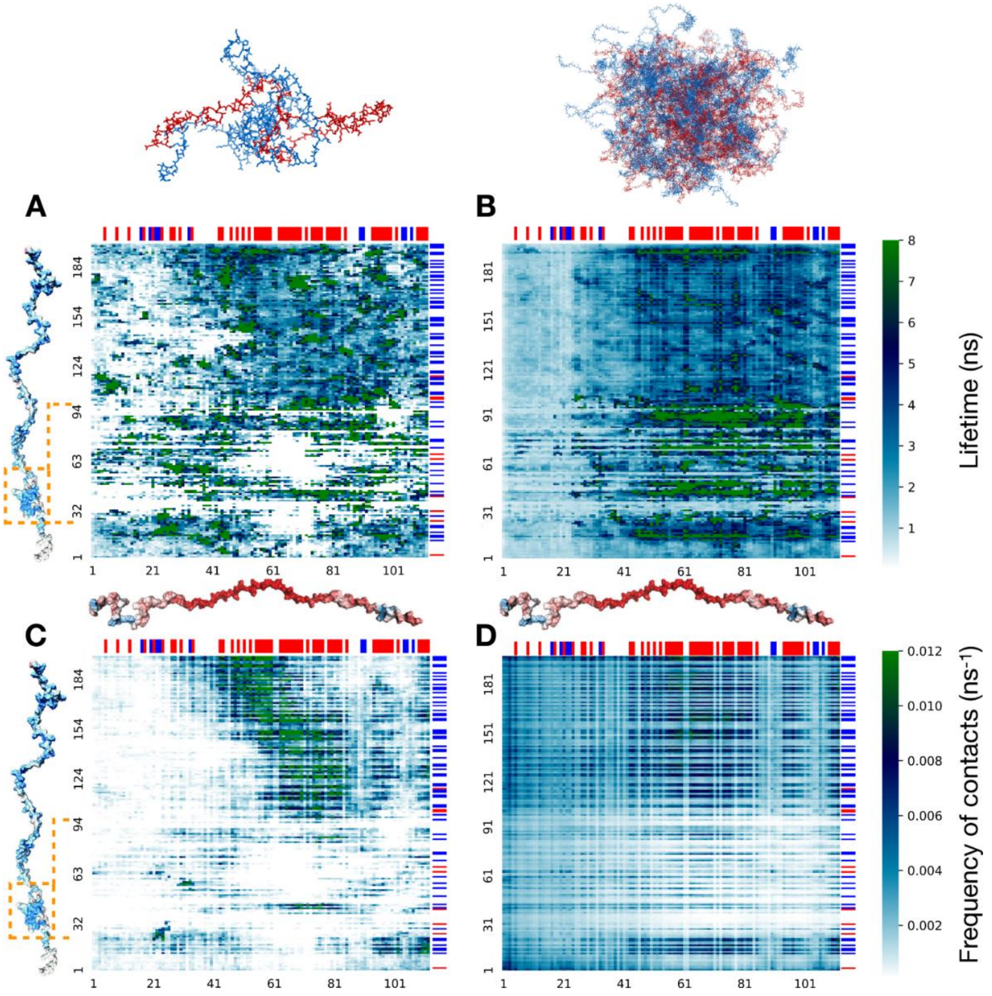


**Figure S7. Contact lifetime heatmaps.** Average lifetime of residue-residue contacts in 6 simulations of the ProTα-H1 dimer (A) and the dense-phase simulation (B). Numbers along the bottom and left denote the residue numbers of ProTα and H1, respectively. Orange rectangles denote the globular domain (GD) of H1 (residues 22 to 96). Frequency of contacts (i.e. the number of newly formed contacts by one ProTα every nanosecond) calculated from dimer and dense phase simulations are shown in (C), and (D), respectively. Blue and red bars at the top and on the right side of the plots denote positively and negatively charged residues of ProTα and H1. In general, the N-terminal part of ProTα makes fewer contacts than the rest of the chain both in the dimer and dense phase simulations (see also Fig 3E), and the lifetime of those contacts is on average shorter, especially in the dense-phase simulation. As is obvious from (D), contacts between oppositely charged residues are most frequent. White regions in A and C correspond to reside-residue combinations that were newer formed during the simulations. White regions are particularly frequent in the GD, since the GD remains folded over the course of dimer simulations (Fig. S13). Some of the GD residues make relatively long-lived contacts, but those contacts are infrequent. In contrast to the dimer simulations, residues of the GD do form contacts with ProTα residues in the dense phase simulation, since a small fraction of partially unfolded GDs are populated (Fig. S13), as expected from the low equilibrium stability of the GD (3,4).

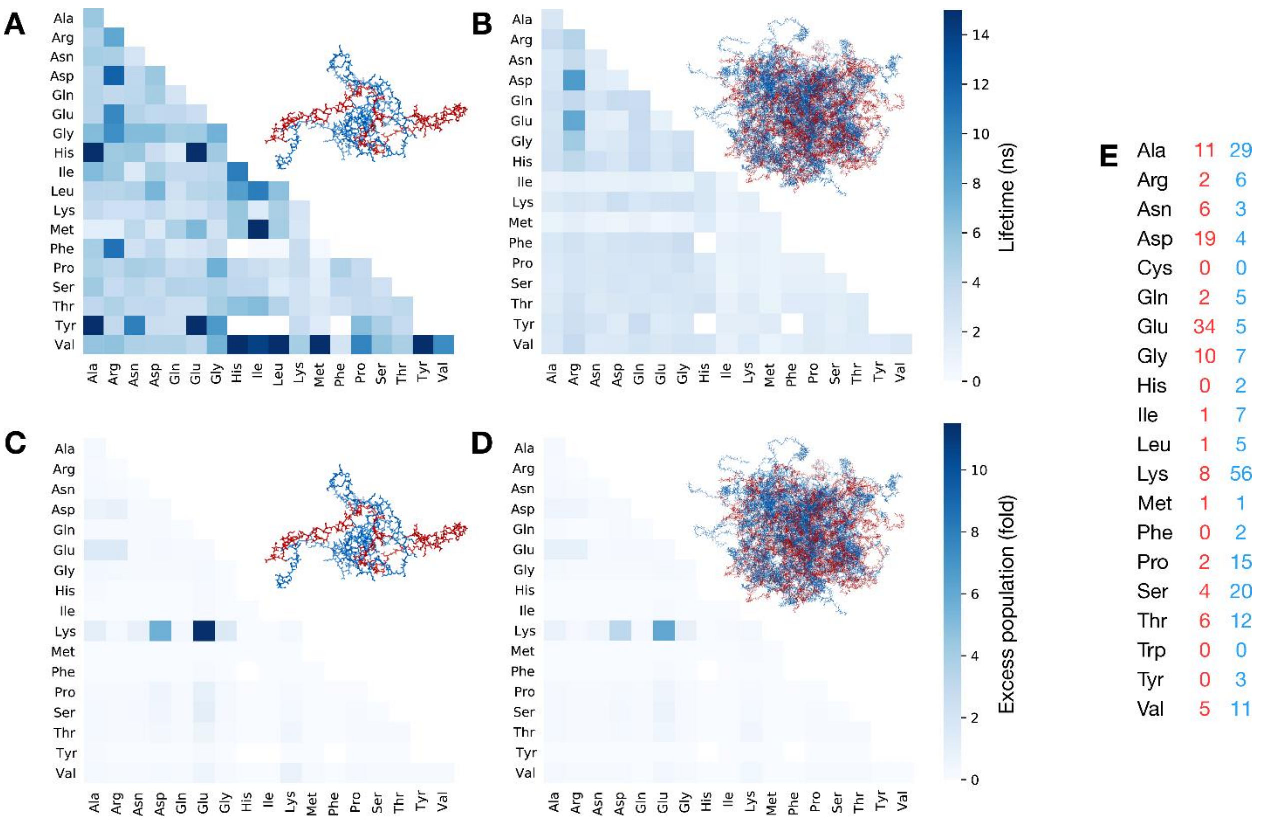


**Figure S8. Residue type-specific contact lifetime heatmaps.** Average lifetimes of residue-residue contacts in the ProTα-H1 dimer (A) and the dense-phase simulations (B) classified by residue types. Excess population of contacts for specific residue pairs in the ProTα-H1 dimer (C) and in the dense-phase simulation (D) (see Methods for details). (E) Numbers of contacts for specific residue types in ProTα (red) and H1 (blue). Residue pairs that are never observed (white squares) and extremely long-lived pairs (dark blue) in (A) correspond to residue types that are infrequent in the ProTα and H1 sequence (compare with (E)). In the dense phase, Arg forms contacts that are on average longer-lived than any other residue (B), in line with the phase separation-promoting role of Arg (5-9). Excess population (see Methods) of contacts for specific residue pairs suggest that the interactions between charged residues are the most favorable interactions both in the dimer and in the dense-phase simulations. Note that the oppositely charged residues Glu (most abundant residue in ProTα) and Lys (most abundant residue in H1) form the largest number of contacts ((C) and (D)) but have lifetimes comparable to other residue pairs ((A) and (B)).

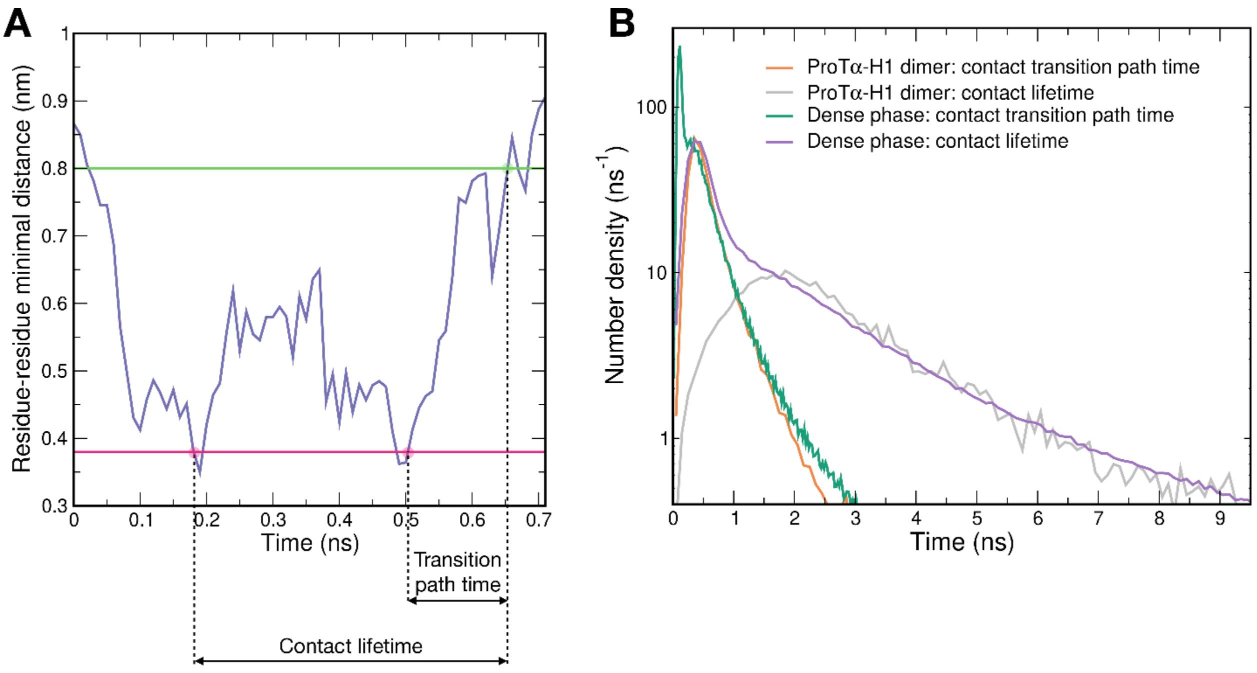


**Figure S9. Estimating the lifetime of non-attractive collisional contacts.** We used the transition path times of residue-residue contacts (A) as an estimate for the lifetime of non-attractive collisional contacts between two residues. While the duration of a contact between two residues was estimated from the point in time when the distance between any two heavy atoms of the two residues falls below 0.38 nm to the time when the distance between any two heavy atoms of those residues reach the 0.80 nm (see Methods), the transition path time of a given contact (for contact breaking) was estimated as a time from the last point in time when the distance between any two heavy atoms of the two residues is below 0.38 nm to the first time it reaches 0.8 nm (A). The timescale expected for non-attractive collisions in the dense-phase simulation (shaded area in Fig. 3F) was estimated as the time that includes 95% of all transition path times in the dense-phase simulation. (B) Comparison between the contact lifetimes and the transition path times in ProTα-H1 dimer and the dense phase. The area under the curves correspond to the total number of contact events per chain per nanosecond.

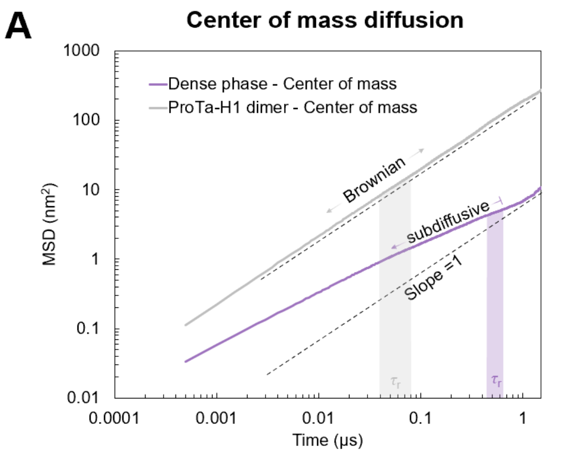


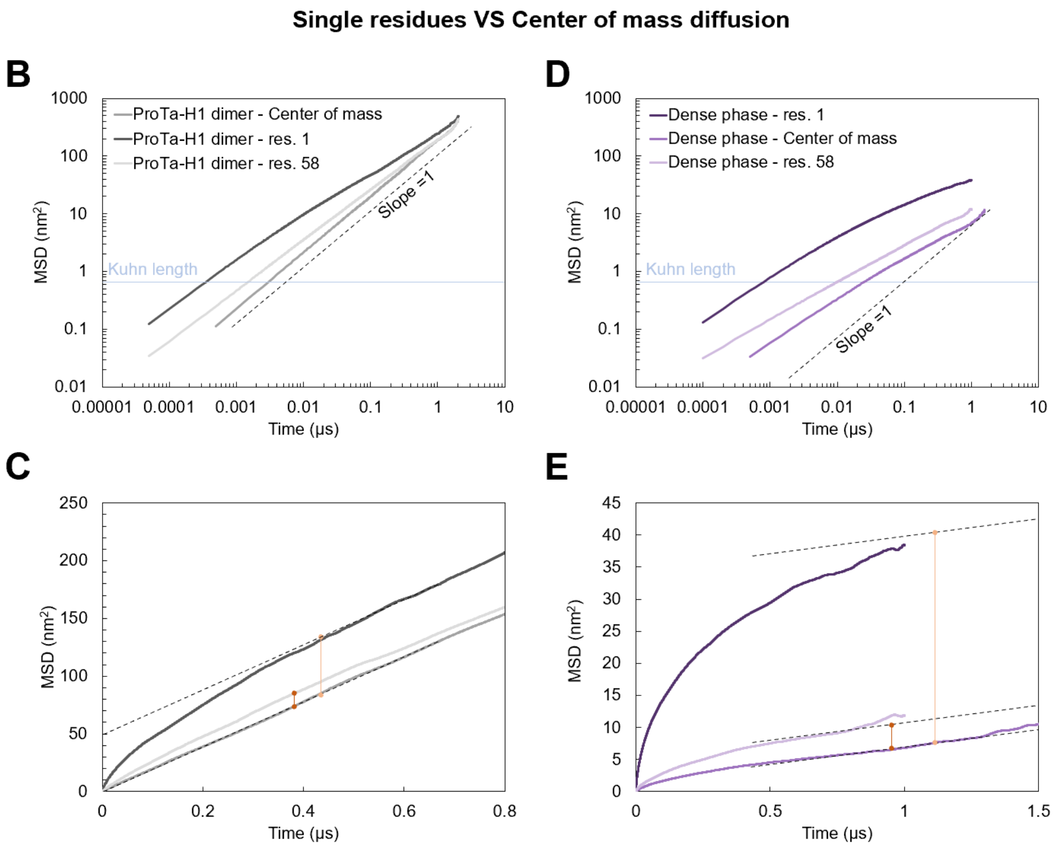


**Figure S10. Mean square displacement (MSD) curves from molecular dynamics simulations reveal subdiffusion.** (A) ProTα center-of-mass diffusion in the dense phase compared to the ProTα-H1 dimer. In the dimer, at all timescales investigated, the diffusion of ProTα is Brownian, whereas in the dense phase, we observe subdiffusive behavior at timescales equal to or shorter than the chain reconfiguration time (shaded bands indicate full-length chain reconfiguration time ± the error), as expected in the presence of cooperative dynamics of the network (10). (B) Comparison between the diffusion of residue 1 of ProTα, the central residue 58, and its center of mass. The residues of an ideal chain are expected to show subdiffusive behavior in a time window between $t_{Kuhn}$, the time a residue needs to diffuse over the Kuhn length of the chain, and the time the entire chain takes to diffuse a distance corresponding to its own size (11), which approximately corresponds to the chain reconfiguration time $\tau_r$ (for a Rouse chain (12)). Below $t_{Kuhn}$, the individual residues are expected to diffuse independently of the chain. Building on the ideal chain model, in the data in Fig. S11, we report the diffusion exponent for timescale below 2 ns (approximately $t_{Kuhn}$), where the single-residue behavior is largely unaffected by the slowdown due to chain reconfiguration. (C) Same data as in (B), but in linear scale to highlight the transition at timescales $>\tau_r$, where the diffusion of the entire chain dominates the diffusion of the individual residues. The yellow and orange vertical lines indicate the MSD travelled by the residue in excess of the MSD of the center of mass of the chain. (D) Analogous to (B) for ProTα in the dense phase. (E) Analogous to (C) for ProTα in the dense phase.

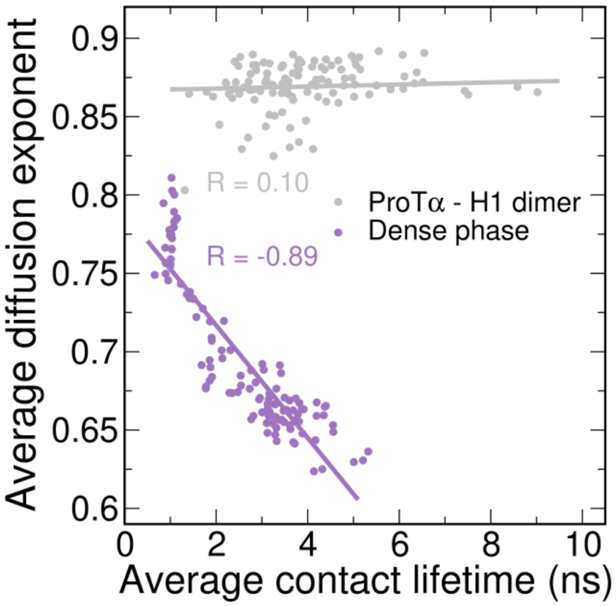


**Figure S11. Diffusive behavior of the individual residues within ProTα.** Diffusion of individual ProTα residues (1-112) is examined in terms of their mean squared displacement, MSD($t$) = $6Dt^{\alpha}$, for timescales shorter than $t_{\text{Kuhn}}$ (see Fig. S10B-E), where $D$ is the diffusion coefficient, $t$ is the lag time, and α = 1 for Brownian diffusion. Diffusion of the residues in the ProTα-H1 dimer is close to Brownian and does not correlate with the average contact lifetime of the corresponding residues, whereas in the dense phase, the diffusion of the residues is more subdiffusive ($\alpha < 1$) and shows a negative correlation with their average contact lifetime. The residues in the dense phase with low average lifetime show less subdiffusive behavior but form the highest number of contacts per unit time (compare with Fig. 3G).

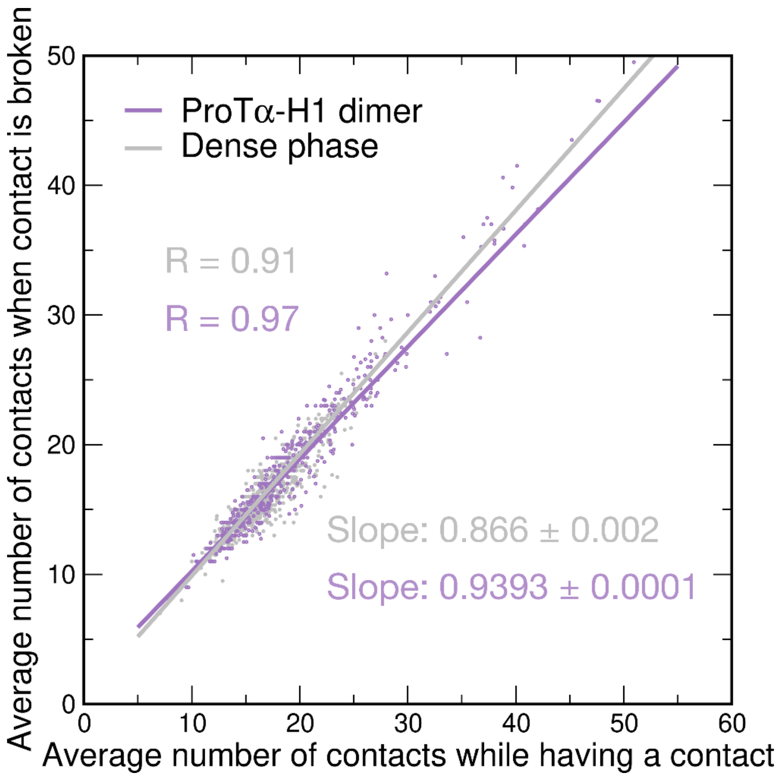


**Figure S12. Competitive substitution between residues.** Average number of contacts at the time when the contact between the two residues is broken plotted as a function of the average number of contacts that those two residues make with other residues during the time being in contact. Given the large number of contact events in the dense phase simulation, only every 20'000$^{th}$ data point is plotted. The definition of a contact is identical to the one described in Methods, but the average number of contacts per residue is larger than the one shown in Fig. 3E since in this case the bonds between neighboring residues were also recorded as contacts. The lower value of the fit slope in the dimer simulation suggests that multiple contacts tend to be broken simultaneously in this case owing to the concerted motions of parts of the protein chains. In contrast, owing to the high local density of potential interaction partners in the dense phase and the competition for contacts, less contacts are broken simultaneously, as the interaction partners are often rapidly substituted (Fig. 3H), resulting in the higher slope in the dense phase simulation.

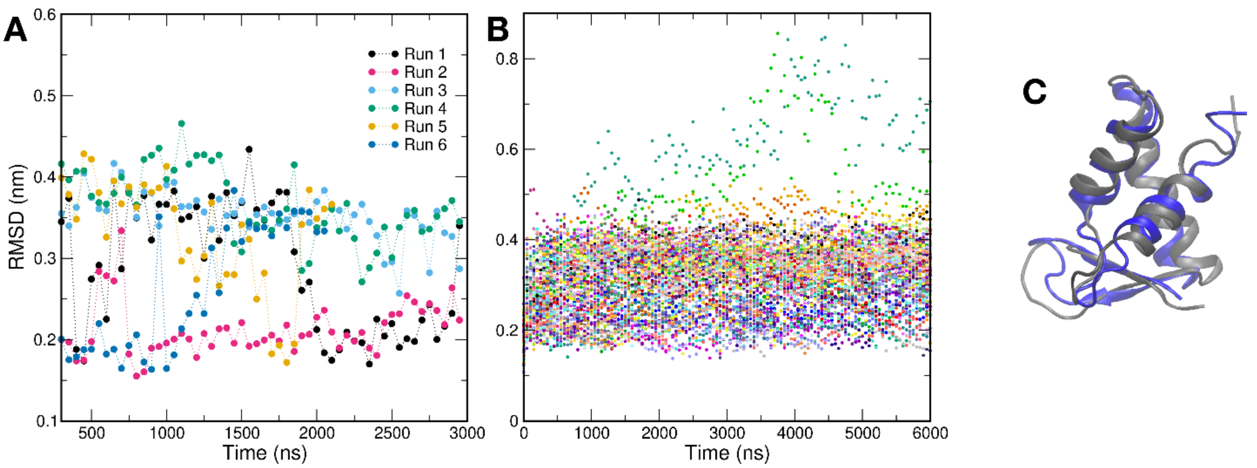


**Figure S13. Stability of H1 globular domains** (GDs), quantified as the backbone RMSD between simulated and experimental structure (PDB 6HQ1) (3), over the course of dimer (A) and dense phase simulations (B). The fraction of partially unfolded structures (< 10% with RMSD > 0.4 nm) is in line with the experimental values previously determined in dilute solution (3). Note that the substantial backbone RMSD of 0.2-0.4 nm even for the folded domain can be attributed to the flexibility of the loops in the structure. (C) Superposition of two structures with RMSD = 0.4 nm, demonstrating that the main difference stems from the flexible parts of GD without global unfolding below this RMSD value.

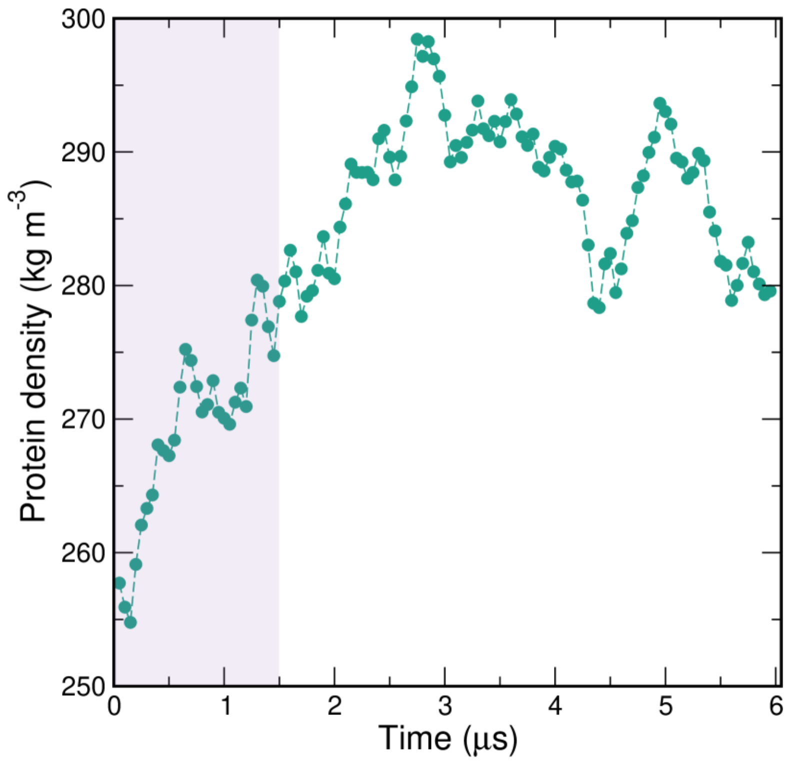


**Figure S14. Equilibration of protein density in the dense-phase simulation.** Protein density in the central part of the slab simulation as a function of time, calculated in 50-ns blocks. The first 1.5 µs of the simulation (shaded band) were considered as equilibration and omitted from further analysis.

| | |
|---|---|
| ProTα<br>(unlabeled) | GPMSDAAVDTSSEITTKDLKEKKEVVEEAENGRDAPANGNANEENGEQEADNE<br>VDEEEEEGGEEEEEEEEGDGEEEDGDEDEEAESATGKRAAEDDEDDDVDTKKQ<br>KTDEDD |
| ProTαN<br>(2C/56C labeled) | G**C**DAAVDTSSEITTKDLKEKKEVVEEAENGRDAPANGNAENEENGEQEADNEVD<br>EE**C**EEGGEEEEEEEEGDGEEEDGDEDEEAESATGKRAAEDDEDDDVDTKKQKT<br>DEDDGA |
| ProTαC<br>(56C/110C<br>labeled) | GPSDAAVDTSSEITTKDLKEKKEVVEEAENGRDAPANGNAENEENGEQEADNEV<br>DEE**C**EEGGEEEEEEEEGDGEEEDGDEDEEAESATGKRAAEDDEDDDVDTKKQ<br>KTDED**C** |
| H1<br>(unlabeled) | TENSTSAPAAKPKRAKASKKSTDHPKYSDMIVAAIQAEKNRAGSSRQSIQKYIKS<br>HYKVGENADSQIKLSIKRLVTTGVLKQTKGVGASGSFRLAKSDEPKKSVAFKKTK<br>KEIKKVATPKKASKPKKAASKAPTKKPKATPVKKAKKKLAATPKKAKKPKTVKAKP<br>VKASKPKKAKPVKPKAKSSAKRAGKKK |

**Table S1.** Amino acid sequences of proteins used. Cys residues introduced for labeling are indicated in bold. Unlabeled ProTα is a variant of human ProTα isoform 2, while ProTα 2C/56C and 56C/110C are variants of isoform 1 (4,13). The isoforms differ by a single Glu at position 39.

**Movies 1 and 2. All-atom explicit-solvent simulation of ProTα-H1 dense phase.** One ProTα chain is highlighted in red, and four interacting H1 chains in different shades of blue. Surrounding ProTα and H1 chains are shown semi-transparently in red and blue, respectively. The image is centered on the center of mass of the highlighted ProTα chain. The total length of the simulation is 6 μs, and the first 1.5 μs were omitted from the analysis; the video is shown at 2 ns per frame. To slightly smooth the motion, a filter with a time constant of 4 ns was applied to all frames. Protein hydrogen atoms, water molecules and ions were omitted for clarity. Movie and 1 and 2 show two different ProTα chains.

**Movie 3. Example of ProTα dynamics at short timescales in the dense phase.** One ProTα chain is highlighted in red, and four interacting H1 chains in different shades of blue. Surrounding ProTα and H1 chains are shown semi-transparently in red and blue, respectively. The image is centered on the center of mass of the highlighted ProTα chain; the video is shown at 100 ps per frame. To slightly smooth the motion, a filter with a time constant of 200 ps was applied to all frames. Protein hydrogen atoms, water molecules and ions were omitted for clarity.
